# Leptin Acts as a Peripheral Tropic Signal to Tune Steroidogenesis

**DOI:** 10.64898/2026.04.13.718290

**Authors:** Jie Sun, Adam Aldahir, Wen-Kan Liu, Calder Ellsworth, Xian-Feng Wang, Hong-Cun Bao, Yi-Chun Huang, Fenglei He, Christoph Heier, Wu-Min Deng

## Abstract

The integration of metabolic status with reproductive and developmental timing is a cornerstone of animal physiology, yet how steroidogenesis rapidly adapts to abrupt environmental changes remains poorly understood. Here, we identify a peptide hormone circuit in which leptin or its analogs act as peripheral tropic signals, directly coupling systemic metabolic state to steroid hormone production through dynamic remodeling of intracellular lipid pools. Mechanistically, spatiotemporal fluctuations in leptin activate JAK/STAT signaling within steroidogenic tissues to reprogram the balance among lipid droplets, cholesteryl esters, and free cholesterol, thereby tuning the amplitude and timing of steroid hormone pulses via hormone-sensitive lipase (Hsl), a rate-limiting determinant of steroidogenic flux. Cross-species analyses in *Drosophila melanogaster*, *Blattella germanica*, and *Mus musculus* suggest a partially conserved leptin-JAK/STAT-lipase axis that functions as a metabolic “sterol rheostat”, enabling rapid modulation of steroidogenesis in response to systemic metabolic and stress cues. These findings reveal a metabolic-endocrine mechanism by which leptin can act directly on steroidogenic organs to regulate hormonal output.

## Introduction

Sexual maturation marks a pivotal developmental transition in which juvenile organisms acquire reproductive competence and adult behaviors. Across metazoans, this transition is driven by steroid hormones that coordinate organismal growth, neural circuit remodeling and reproductive activation ^1, 2^. In mammals, steroid hormones reorganize brain circuitry during the perinatal and pubertal periods and subsequently promote sexual drive in adulthood ^3, 4^. In insects, pulses of the steroid hormone ecdysone produced by the prothoracic gland (PG) similarly orchestrate major developmental transitions, including metamorphosis, in a process functionally analogous to mammalian puberty ^5, 6^. Despite this deep evolutionary conservation, how steroidogenic tissues rapidly adjust hormone output to match changing metabolic and developmental demands remains poorly understood.

Steroidogenesis requires precise coordination of lipid and cholesterol metabolism in endocrine cells ^7^. Because steroid hormones themselves are poorly stored, timely secretion is dependent on the rapid mobilization of cholesteryl ester (CE) ^8^, which buffers excess free cholesterol (FC) and provides a readily accessible precursor pool ^9^. Together with triacylglycerol (TG), CEs are packaged into intracellular lipid droplets (LDs) ^10^, conserved organelles that buffer fluctuations in lipid availability ^11^. Upon physiological demand for steroid hormones, cells recruit lipases to the LD surface to catalyze ester hydrolysis, thereby releasing FC for mitochondrial steroidogenesis ^9, 12–14^. Accordingly, efficient developmental transitions likely require tightly regulated LD remodeling. However, whether LDs and CEs undergo rapid, signal-dependent reorganization in response to systemic developmental or nutritional cues remains unknown.

In steroidogenic cells, hormonal output is classically governed by gonadotropic peptide signals ^15, 16^. Traditionally viewed as endocrine factors mediating long-range communication between distant tissues, peptide hormones are now increasingly recognized as being synthesized within developing tissues, where they contribute to the maturation of the hypothalamic-pituitary-gonadal (HPG) axis during puberty ^17, 18^. Among systemic metabolic cues, leptin, produced primarily by adipocytes and enterocytes ^19, 20^, has emerged as a conserved integrator of energy balance and reproductive capacity across species ^21, 22^. Leptin facilitates the pubertal transition and reproductive homeostasis by stimulating hypothalamic gonadotropin-releasing hormone (GnRH) neurons to drive gonadotropin secretion ^21, 23, 24^, and circulating leptin levels dynamically reflect nutritional status ^25^. Yet physiological and clinical observations, including the paradoxical association of hyperleptinemia with hypogonadism in obesity and inflammatory states ^26, 27^, suggest that additional leptin-responsive mechanisms remain unresolved. In particular, whether leptin can act directly on steroidogenic tissues, independent of the central nervous system, to rapidly couple organismal metabolic state with steroid output is unknown.

To address this question, we performed continuous spatiotemporal tracking of the LD-CE-steroid flux in endocrine cells from representative holometabolous and hemimetabolous insect species and mouse. Our findings uncover a previously unrecognized metabolic-endocrine pathway that enables rapid, context-dependent tuning of steroidogenesis. We show that steroidogenic cells themselves function as direct systemic sensors of leptin, remodeling intracellular lipid substrate availability to control circulating steroid levels, thereby ensuring robust and timely sexual maturation during the juvenile-to-adult transition.

## Results

### Dynamic remodeling of the LD-CE pool in steroidogenic cells during development

LDs are highly dynamic organelles that serve as reservoirs for the sequestered storage of CEs, the indispensable metabolic substrates for steroidogenesis. To determine whether LDs participate in developmental timing, we investigated LD dynamics in steroidogenic cells from multiple model systems, beginning with mouse testicular Leydig cells. These cells are primarily responsible for the production and secretion of the steroid hormone testosterone, a key regulator of male sexual development. Developmental profiling revealed pronounced age-dependent remodeling of the LD pool: Bodipy staining showed that immature Leydig cells (3 weeks old) were nearly devoid of LDs, whereas LDs progressively accumulated during sexual maturation (5-9 weeks old) and reached robust steady-state levels in fully mature cells at 6 months of age (Fig. 1A, B).

**Figure 1.**
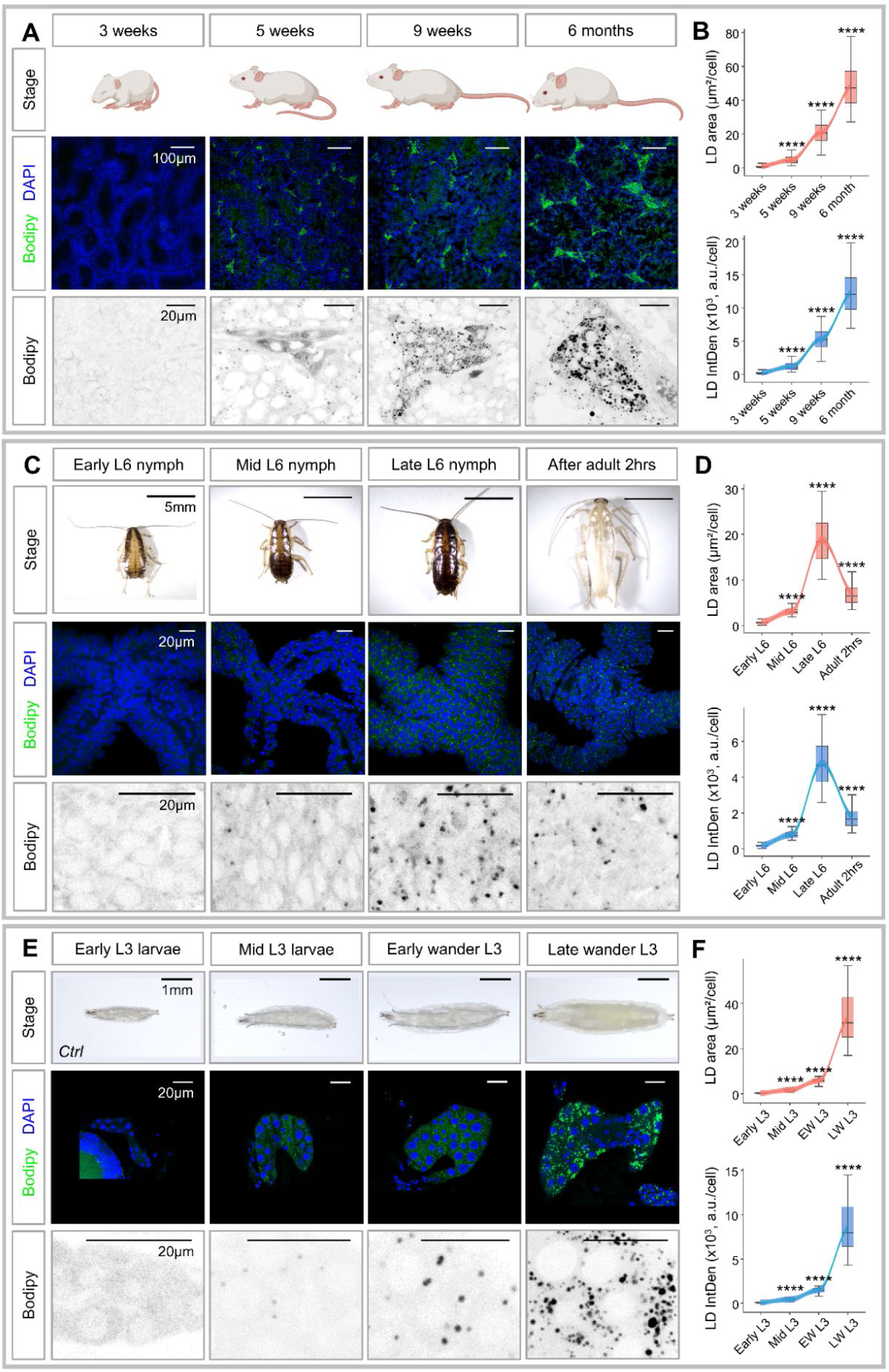
LD pool remodeling in steroidogenic tissues in response to the demands of sexual development. (A) Confocal microscopy images of LD dynamics in male mice Leydig cells across ages: 3 weeks, 5 weeks, 9 weeks, and 6 months. Top panels: Whole-animal diagrams. Middle panels: Low-magnification confocal sections of Bodipy (green)/DAPI (blue) staining (scale bar, 100 µm). Bottom panels: High-magnification views of grayscale Bodipy images (scale bar, 20 µm). (B) Quantification of LD area and fluorescence intensity during sexual development in mouse. (C) Confocal microscopy images showing LD dynamics stained with Bodipy (green) and nuclei with DAPI (blue) in the PG across developmental stages in *Blattella germanica* (early L6 nymph, mid-L6 nymph, late L6 nymph, and 2 hours after eclosion adult). From top to bottom panel: Whole-animal images (scale bar, 5 mm); low-magnification confocal sections of Bodipy/DAPI staining (scale bar, 20 µm); high-magnification views of representative regions indicated in the middle panels (scale bar, 20 µm). (D) Quantification of LD area and fluorescence intensity during sexual development in *Blattella germanica*. (E) Confocal microscopy images showing LD dynamics stained with Bodipy (green) and nuclei with DAPI (blue) in PG cells across developmental stages in *Drosophila* (early L3, mid-L3, early wandering L3, and late wandering L3) under control conditions (*w1118*). From top to bottom: whole-larva images (scale bar, 1 mm); low-magnification confocal sections of Bodipy/DAPI staining (scale bar, 20 µm); high-magnification views of representative regions indicated in the middle panels (scale bar, 20 µm). (F) Quantification of LD area and fluorescence intensity over developmental time shows a significant increase in control (*w1118*). *P* values are calculated using one-way analysis of variance (ANOVA) followed by Holm-Sidak multiple comparisons. Asterisks illustrate statistically significant differences between conditions, ****p<0.0001.

Unlike mammals, insects lack the capacity for *de novo* cholesterol biosynthesis and instead rely on dietary sterols that are transported through the hemolymph by lipophorin and stored primarily in the fat body and steroidogenic tissues ^28–30^, providing physiologically simplified models for investigating the dynamic regulation of LD-CE homeostasis. In the hemimetabolous cockroach *Blattella germanica*, Bodipy staining revealed striking developmental remodeling of LD pools in the PG from the final nymphal instar (L6) to the adult stage. LDs were nearly absent during the early and mid-L6 nymphs. As animals progressed into late L6, LDs expanded substantially in both size and abundance, eventually filling much of the PG cytoplasm (Fig. 1C). Following adult eclosion, these LDs were progressively hydrolyzed (Fig. 1C). Quantitative analysis confirmed that the marked expansion of LD size and fluorescence intensity coincided with the onset of sexual maturation (Fig. 1D).

To dissect the regulatory mechanism underlying LD remodeling in a genetically tractable model, we next turned to the holometabolous model insect *Drosophila melanogaster*. Consistent with other systems, LD accumulation in the *Drosophila* PG was highly dynamic. LDs were sparse during early- and mid-L3 stages but progressively expanded during late L3, eventually occupying most of the PG cytoplasm by the wandering stage prior to pupariation (Fig. 1E). Quantification revealed a significant increase in LD accumulation between 72 and 130 hours after egg deposition (AED), precisely coincident with the steroidogenic maturation window (Fig. 1F). Furthermore, LDs colocalized robustly with NBD-cholesterol (22-NBD-cholesterol) in PG cells, indicating that extracellularly acquired FCs were preferentially sequestered into LDs for downstream steroid hormone biosynthesis (Fig. S1A, B). To resolve temporal behavior, we performed real-time imaging of LDs in live PG cells. Time-lapse microscopy over an 8-hour window revealed continuous, progressive LD remodeling synchronized with endocrine transitions (Fig. S1C; Supplementary Movies S1, S2).

Together, these findings demonstrate that the cholesterol-storing LD pools in steroidogenic cells are tightly and dynamically regulated during development, exhibiting a conserved remodeling program during sexual maturation that poises cells for stage-appropriate steroidogenic output across species.

### JAK/STAT signaling governs LD dynamics in steroidogenic cells

To identify the molecular drivers of developmental LD remodeling, we performed transcriptomic profiling of the PGs at mid-L3 and 18 hours later in late-L3 larvae. Differential gene set enrichment analysis (GSEA) revealed a progressive downregulation of JAK/STAT signaling components, including the cytokine receptor *domeless* (*dome*), the Janus kinase *hopscotch* (*hop*), and the downstream target gene *Socs36E*, coincident with the onset of sexual maturation. Conversely, negative regulators of the pathway (*Su(var)2-10*, *Ptp61F*, and *ken*) showed increased expression during this window (Fig. S2A, B). These transcriptomic trends were corroborated by the 10xSTAT::GFP reporter ^31^, which showed robust activation in mid-L3 PG cells that declined toward late L3 (Fig. 2C). Time-lapse imaging further revealed dynamic STAT activity over a 6-hour period during the mid-to-late L3 stage, with GFP intensity inversely correlated with LD abundance, such that peaks in GFP signal consistently coincided with periods of low LD occupancy (Fig. 2A, B; Supplementary Movies S3, S4). Moreover, pharmacological inhibition of JAK/STAT signaling using AG490 ^32^ during early L3 triggered premature LD expansion, with a pronounced increase in LD area (Fig. 2D, E).

**Figure 2.**
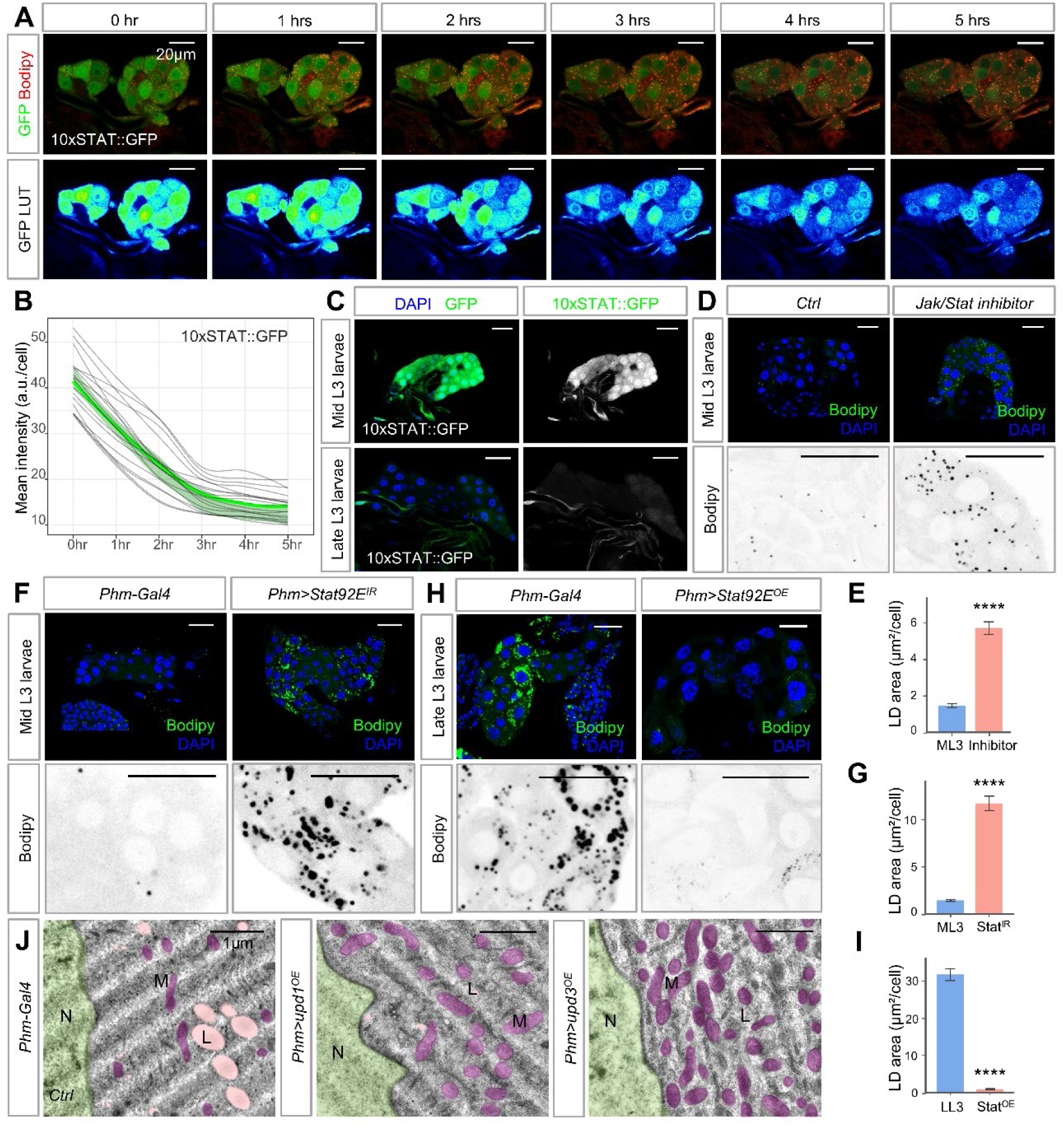
JAK/STAT signaling modulates the LD reservoir in the PG during development. (A) Time-lapse imaging over 6 hours of PG cells in 10xSTAT::GFP larvae. Fluorescence intensity in the GFP channel indicates STAT activity, and Bodipy (red) labels LDs. Corresponding Bodipy images are shown below each panel with LUT representation. (B) Quantitative analysis shows the Fluorescence (GFP channel) intensity (JAK/STAT activity) in PG cells over the 5-hour imaging period. (C) JAK/STAT signaling activated level in the PG cells show by 10xSTAT::GFP (green) with DAPI (blue) marking nuclei at mid- and late-L3 instar larval. (D) Representative Bodipy staining (green) and quantification of LD area (E) in PG cells following 24h treatment with the JAK/STAT inhibitor compared to control at the mid-L3 stage. (F) At mid-L3 larvae, Bodipy (green) staining of LDs in the PG cell under control (*Phm-Gal4*) and *Stat92E RNAi* at mid-L3 larvae. (H) At late-L3 larvae, LDs (green) reduced in Stat92E overexpression conditions, compared to controls. In both (F) and (H), top panels show low magnification of confocal section, and bottom panels show high-magnification view of representative areas in top panels. (G and I) Quantification of LD area per cell for genotypes in (F) and (H). (J) TEM images showing the distribution of LD (L), nuclei (N), and mitochondria (M) in the PG of control (*Phm-Gal4*) larvae and larvae with PG-specific overexpression of Upd1 or Upd3. Scale bars, 1 μm.

To establish the functional requirement of JAK/STAT signaling in LD remodeling, we performed stage-specific genetic manipulations in the PG of *Drosophila*. At mid-L3, PG-specific knockdown of *hop* or *Stat92E*, or expression of a dominant-negative *dome* receptor (*dome^DN^*) via *phm-Gal4* ^33^, significantly increased LD size relative to controls (Fig. 2F, G and Fig. S2C). Conversely, overexpression of Stat92E or the ligands Upd1 and Upd3 in late-L3 PGs suppressed LD accumulation (Fig. 2H, I and Fig. S2D), indicating that JAK/STAT activity restricts LD formation. Consistent with these observations, transmission electron microscopy (TEM) provided ultrastructural confirmation, revealing that the electron-dense LDs characteristic of control PG cells was markedly reduced upon Upd1 or Upd3 overexpression (Fig. 2J). Together, these data establish JAK/STAT signaling as a developmental brake whose progressive decline permits stage-specific LD accumulation in the PG.

### Fru is autonomously regulated by the JAK/STAT pathway to control LD remodeling

To explore candidate downstream transcription effector, we examined the zinc-finger protein Fruitless (Fru), previously implicated in lipid metabolism in oenocytes, the insect hepatocyte-like cells ^34^. Using the *fru-NP21-Gal4> UAS-GFP* reporter ^35^, we found that *fru*, is expressed in the PG (Fig. S3A). Immunofluorescence further revealed that Fru protein levels declined progressively as LD abundance increased from early-to late-L3 stages (Fig. 3A, B). Consistent with this pattern, RNA-seq analysis confirmed stage-dependent regulation of *fru*, with transcript levels in mid-L3 PGs significantly higher than those detected 18 hours later (Fig. 3D). Moreover, quantitative correlation analysis between LD distribution and Fru expression revealed a pronounced inverse relationship (Fig. 3C). Functionally, PG-specific knockdown of *fru* induced premature LD accumulation at mid-L3 (Fig. 3E, F; Fig. S3B, C). In contrast, Fru overexpression at late L3 reduced LD size and density (Fig. 3G; Fig. S3B), a phenotype reproduced using an independent PG driver (*Spok-Gal4*^36^) (Fig. S3D). TEM analysis further confirmed that Fru antagonizes lipid storage within the PG (Fig. 3H).

**Figure 3.**
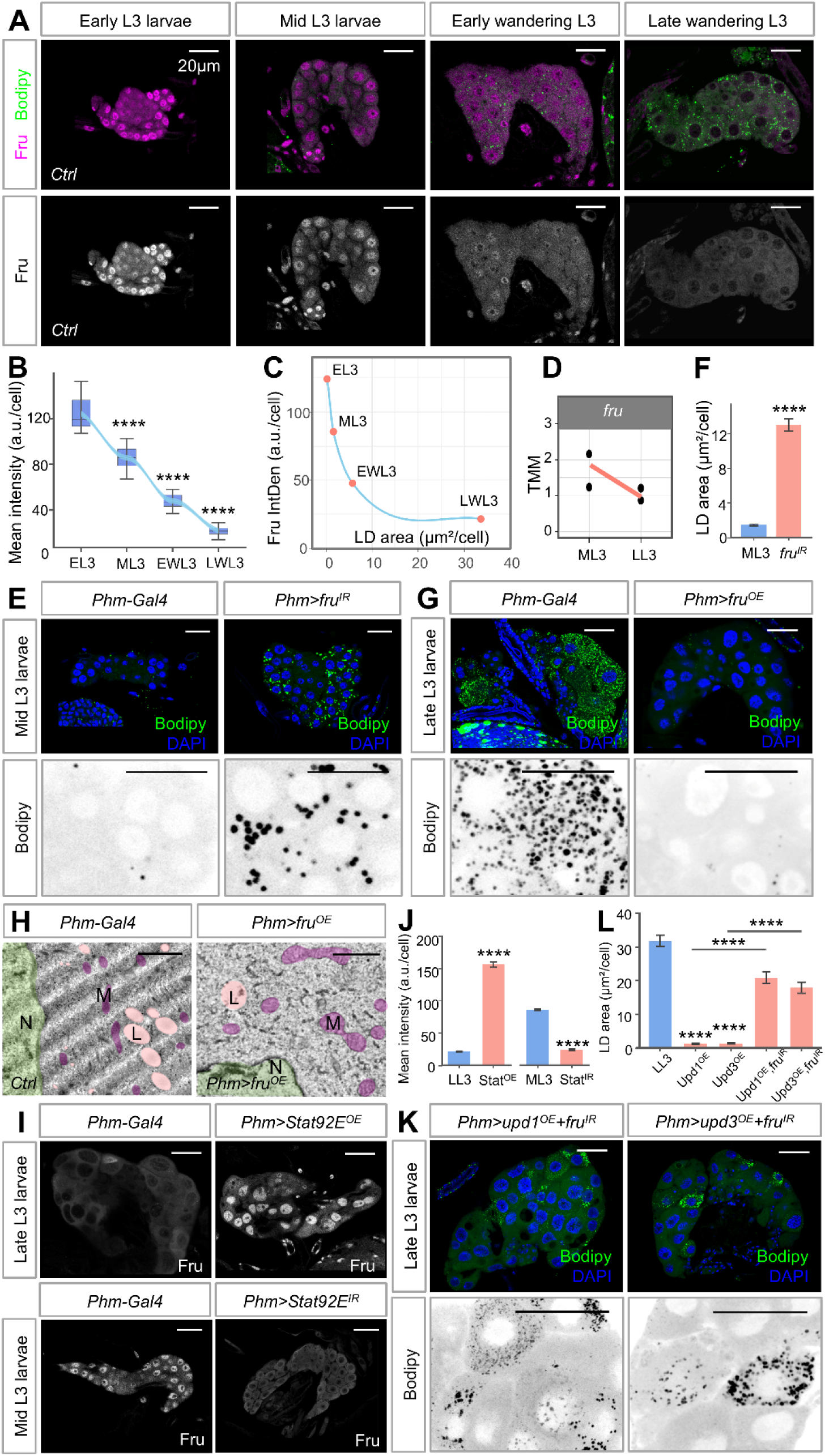
Fru acts downstream of JAK/STAT to control LD remodeling in the PG. **(A)** Immunofluorescence staining show Fru expression pattern (magenta) and LDs (Bodipy, green) in the PG across developmental stages (early L3, mid-L3, early wandering L3, and late wandering L3) under control (*w1118*). (B) Quantitative analysis of Fru fluorescence intensity and correlation between Fru intensity and LD area across developmental stages (C). (D) RNAseq analysis shows *fru* mRNA levels in mid-L3 versus late-L3 larvae (p<0.05). (E) Representative Bodipy staining (green) and quantification of LD area (F) in PG cells under control (*Phm-Gal4*) and *fru RNAi* at mid-L3 larvae. (G) Representative images of Bodipy (green) and DAPI (blue) in PG cells under control and *phm>* (PG-specific) with Fru overexpression at late L3 larvae. Top panels show low-magnification confocal sections; bottom panels present high-magnification views of representative regions indicated in the top panels. (H) TEM images of LD (L), nuclei (N), and mitochondria (M) in the PG of control (*Phm-Gal4*) larvae and larvae with PG-specific overexpression of Fru. Scale bars, 1 μ m. (I) Immunofluorescence images and quantification (J) of Fru expression in the PG under PG-specific control (*phm>*), showing Stat92E overexpression in late-L3 larvae and *Stat92E* knockdown in mid-L3 larvae. (K) Representative images and quantification (L) showing the LDs (Bodipy, green) in the PG of late-L3 larvae, under PG-specific Fru knockdown in the background of JAK/STAT activation (Upd1or Upd3 overexpression). For all graphs, statistical significance was followed by one-way ANOVA or Student’s T-test. ****p<0.0001. Scale bars in confocal images represent 20μm unless otherwise noted.

Given the strong concordance between Fru expression and JAK/STAT activity during L3 (Fig. S4A), we tested whether these two factors functionally interact. Upstream activation of the JAK/STAT pathway by overexpression of Upd1, Upd2, Upd3, Hop, or Stat92E robustly elevated Fru levels in late-L3 PGs (Fig. 3I, J; Fig. S4D). Conversely, inhibition of the pathway via *dome^DN^*, *hop RNAi* or *Stat92E RNAi* significantly reduced Fru expression in mid-L3 PGs (Fig. 3I, J; Fig. S4E). Notably, Fru overexpression did not activate the JAK/STAT signaling nor induce *upd3* transcription (Fig. S4B, C), placing Fru downstream of the pathway. Importantly, co-expression of *fru RNAi* with Upd1 or Upd3 overexpression rescued the LD phenotypes induced by JAK/STAT activation (Fig. 3K, L), whereas simultaneous overexpression of Fru and knockdown of *upd3* failed to restore LD accumulation (Fig. S4F). These epistasis experiments indicate that Fru is required for JAK/STAT-mediated control of LD remodeling.

We next asked whether the JAK/STAT-Fru axis coordinates developmental progression by regulating LD dynamics in the PG. PG-specific knockdown of *fru* markedly accelerated the larval-to-pupal transition, whereas overexpression of Fru isoforms (FruB or FruC) caused severe developmental arrest at the L1 stage (Fig. S5A, B), indicating that precise Fru dosage is essential for developmental timing. Direct measurement of ecdysteroid titers showed that two independent *fru* knockdowns significantly elevated hormone levels during the early- and late-wandering L3 stages (Fig. S5C). Because constitutive Fru overexpression caused severe developmental arrest at L1, we used *tub-GAL80^ts^*, a temperature-sensitive GAL4 inhibitor ^37^, to induce Fru expression from early L3 stage. Under these conditions, ecdysteroid titers were significantly reduced in Fru-overexpressing larvae relative to controls during both early and late wandering L3 stages (Fig. S5C), demonstrating that Fru negatively regulates steroid output from the PG. Functionally, dietary supplementation with 20-hydroxyecdysone (20E) fully restored pupariation and larval mouth-hook morphology (Fig. S5D-F). Together with prior evidence linking JAK/STAT signaling to developmental transitions ^38^, these data demonstrate that the JAK/STAT pathway cell-autonomously regulates LD remodeling through its downstream effector Fru in *Drosophila* PG, thereby dynamically controlling sterol substrate availability and steroid hormone output, and ensuring robust coordination of sexual maturation and developmental timing.

### Hormone-sensitive lipase: A metabolic bottleneck in JAK/STAT-Fru-directed LD formation

To determine how lipid homeostasis is regulated in the PG, we performed transcriptional profiling following Fru perturbation. Gene Ontology (GO) enrichment analysis of Fru-overexpressing PGs revealed that downregulated genes were enriched in developmental and metabolic programs, including fatty acid metabolism, fatty acyl-CoA metabolism, and chitin metabolism (Fig. 4A). GSEA further indicated coordinated but modest repression of metabolic pathways, with reduced expression of genes involved in fatty acid metabolism and lipid biosynthesis (Fig. 4B), with LD regulatory proteins being particularly affected (Fig. 4C). Time-course transcriptomic analysis identified the lipolytic regulator Hormone-sensitive lipase (Hsl) as prominently downregulated during normal PG maturation (Fig. 4D), mirroring Fru dynamics and the progressive expansion of the LD pool.

**Figure 4.**
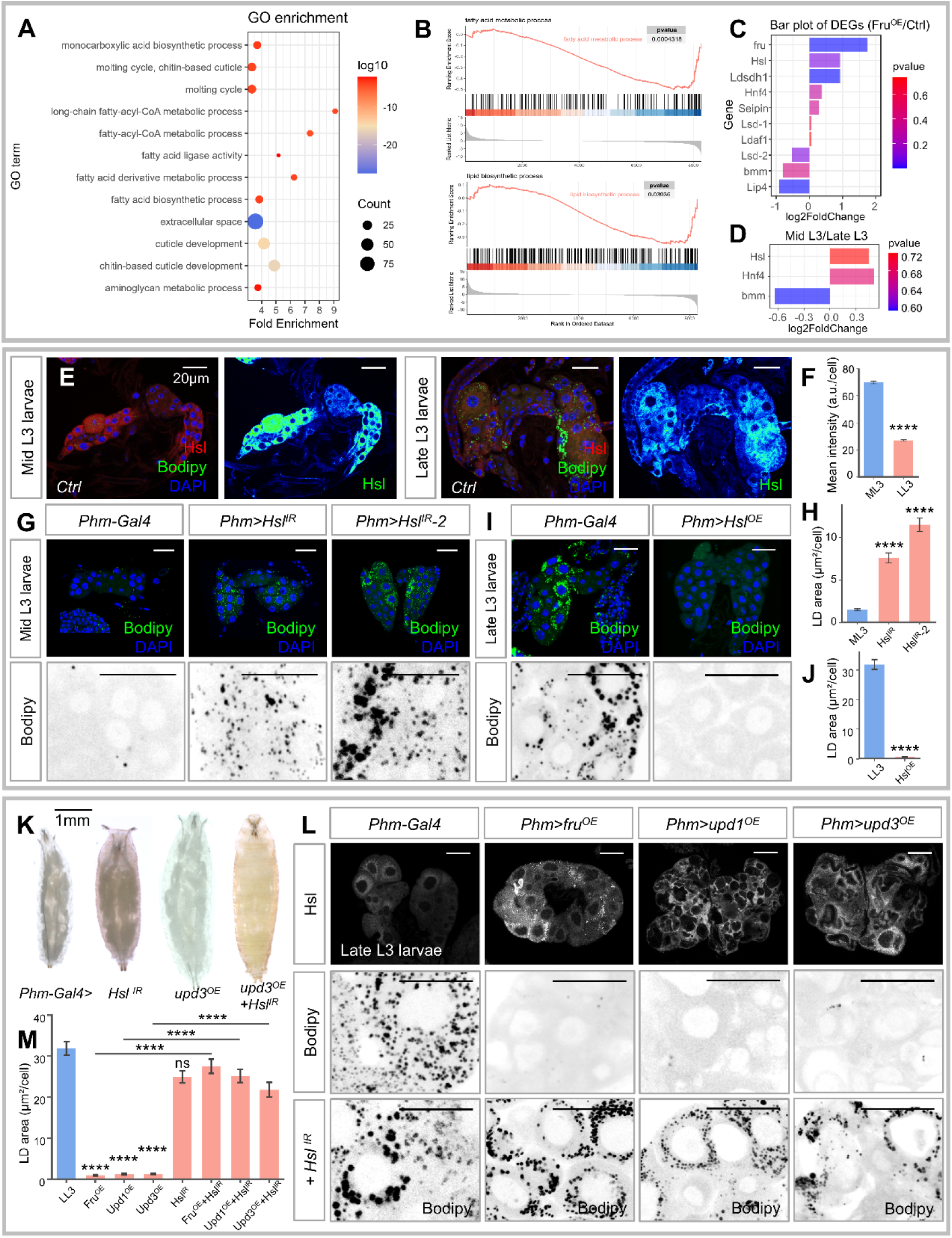
The JAK/STAT-Fru axis regulates Hsl expression to modulate lipolysis in the PG. (A) Gene Ontology (GO) enrichment analysis of differentially expressed genes (DEGs) in PGs upon Fru overexpression. The *y* axis indicates the pathway name, and the *x* axis indicates the fold change. (B) Gene Set Enrichment Analysis (GSEA) showing enrichment of the fatty acid metabolic process and lipid biosynthetic process gene sets in *Phm>fruB^OE^*compared with control (*phm-GAL4*). (C) Bar plots of DEGs showing changes in lipid metabolism related genes in the PG with *fru* knockdown. (D) Bar plot of DEGs shows the change of lipolytic regulators in mid-L3 versus late-L3 PG cells. Representative images (E) and quantification (F) of Hsl staining in the PG of control (*w1118*) at the mid-L3 and late-L3 larvae. Representative images of Bodipy (green) and DAPI (blue) in PG cells under control of *phm>* (PG-specific) with Hsl knockdown (*Hsl-RNAi-1* and *Hsl-RNAi-2*) at mid-L3 larvae (G), and with Hsl overexpression at late-L3 larvae (I). Top panels show low-magnification confocal sections; bottom panels present high-magnification views of representative regions indicated in the top panels. (H and J) Graphs showing quantification of LD area in the PG based on images (represented in G and I) normalized to controls. (K) Larvae and prepupal phenotypes at 120 h after egg laying, including control (*phm>*), PG-specific knockdown of *Hsl* and the rescue of the Upd3^OE^ phenotype by co-knockdown of *Hsl*. (L) Representative images showing Fru and JAK/STAT activation (*Phm>fru^OE^*, *upd1^OE^* and *upd3^OE^*) upregulate Hsl expression in the PG at late L3 larvae (top panels), and concomitant *Hsl* knockdown increases LD accumulation (bottom panels). (M) Quantification of LD area per cell based on the genotypes shown in (L). Data are represented as means ± SEM. Statistical significance was determined by one-way ANOVA followed by multiple comparisons. ****p < 0.0001; ns, not significant. Scale bars represent 20μm for confocal images and 1 mm for larvae.

Direct examination of Hsl protein confirmed this developmental regulation. Immunofluorescence analysis showed that Hsl levels declined from mid- to late-L3 (Fig. 4E, F). Functional perturbations established Hsl as a rate-limiting regulator of LD turnover: PG-specific knockdown of *Hsl* using two independent RNAi lines markedly increased LD accumulation at mid-L3, whereas Hsl overexpression suppressed LD size and density at late L3 (Fig. 4G-J). Notably, premature LD accumulation caused by Hsl depletion accelerated the larval-to-pupal transition, indicating that precise control of Hsl-mediated lipolysis is required for proper developmental timing (Fig. 4K and Fig. S6A).

To test whether Hsl functions downstream of the JAK/STAT-Fru axis, immunostaining with anti-Hsl antibodies showed that Fru overexpression or ectopic JAK/STAT activation markedly increased Hsl levels in PG cells (Fig. 4L and Fig. S6C, D). Consistently, a GFP-tagged endogenous Hsl protein trap exhibited enhanced signal upon Fru overexpression (Fig. S6B), supporting transcriptional regulation of Hsl by Fru. Furthermore, *Hsl* knockdown under conditions of hyperactivated JAK/STAT-Fru axis restored LD accumulation in late-L3 PGs (Fig. 4L, M and Fig. S6C). Functionally, *Hsl* depletion also rescued the eclosion defects induced by Upd3 overexpression (Fig. 4K). Together, these data identify Hsl as a critical downstream effector of the JAK/STAT-Fru axis and demonstrate that lipolytic control in the PG constitutes a rate-limiting step in LD remodeling and developmental progression.

Because Hsl catalyzes the conversion of CE to FC ^39^, we next asked whether sustained activation of the JAK/STAT signaling or Fru overexpression produces pathological FC overload. Filipin staining, which specifically detects unesterified cholesterol ^40^, revealed that activation of JAK/STAT signaling (via Upd3 or Stat92E overexpression) or Fru overexpression induces excess intracellular FC deposition (Fig. 5A). Ultrastructural TEM analysis further showed that accumulated FC formed intracellular cholesterol monohydrate crystals (Fig. 5B). Because cholesterol crystallization is associated with fibrosis and calcification ^41, 42^, we performed wheat germ agglutinin (WGA) and Alizarin Red S staining, which label glycoprotein-rich matrix components and calcium deposition, respectively ^43^, confirming pronounced fibrosis-like matrix accumulation and calcification in Fru-overexpressing PG cells (Fig. 5C, D).

**Figure 5.**
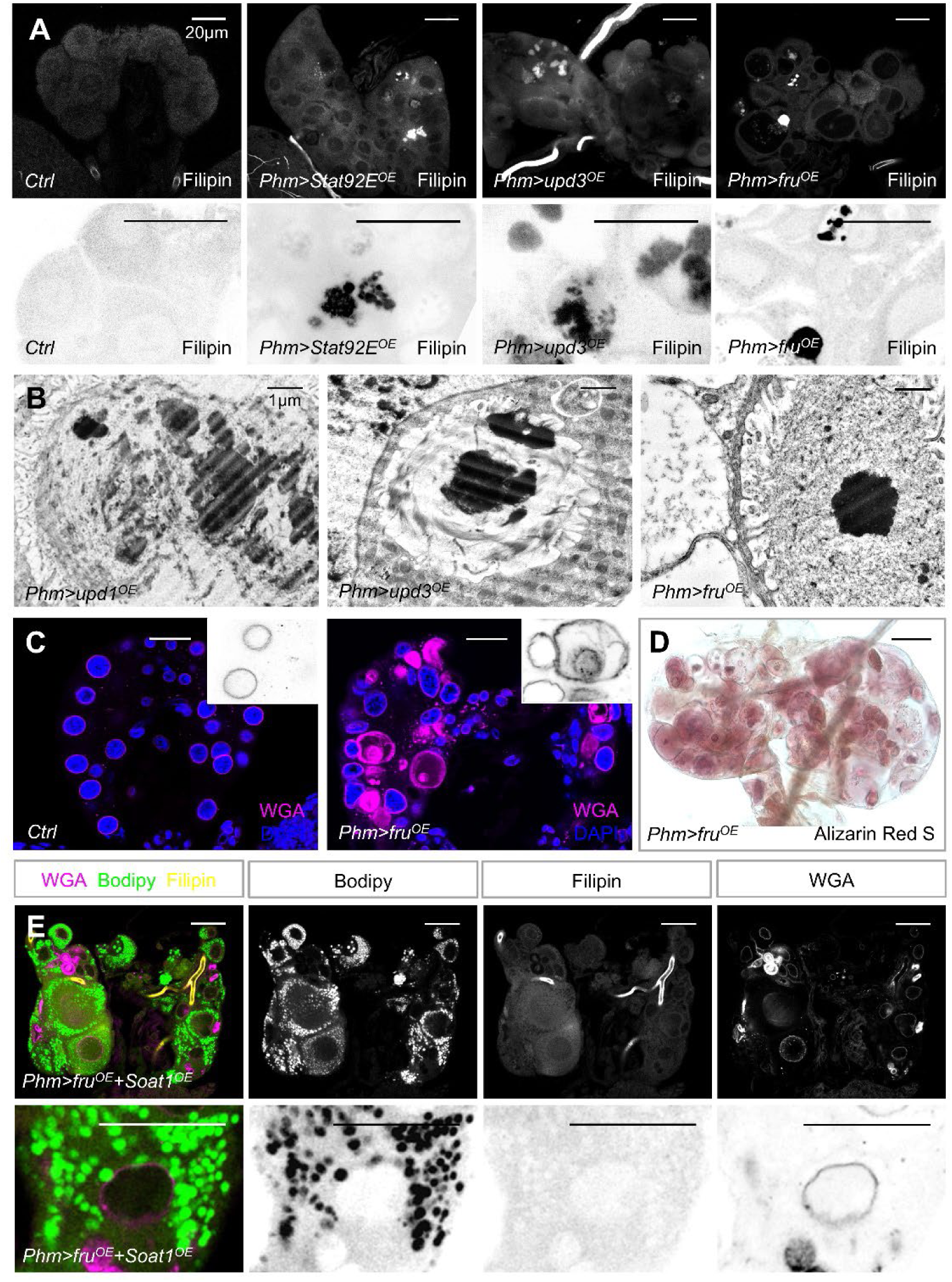
Sustained activation of the JAK/STAT-Fru-Hsl axis induces intracellular cholesterol crystallization. (A) Representative images of Filipin staining in PG cells of control (*phm-GAL4*) and genotypes with overexpression of Stat92E, Upd3, and Fru. Lower panels show high-magnification inverted gray-scale images highlighting the presence of dense cholesterol aggregates (black) in the overexpressed groups. (B) Representative TEM micrographs show cholesterol crystals from PGs of *phm>upd1^OE^*, *phm>upd3^OE^*, and *phm>fru^OE^* larvae. Scale bar: 1 μ m. (C) Immunofluorescence staining of PG cells stained with WGA (magenta) and DAPI (blue) under control (*phm-GAL4*) and *phm>* (PG-specific) with Fru overexpression at late L3 larvae. (D) Representative image of the PG from *phm>fru^OE^*stained with Alizarin Red S. (E) Multichannel fluorescence imaging of PGs with co-overexpression of Fru and Soat1. Top panels: Representative images showing the distribution of WGA (magenta), Bodipy (green), and Filipin (yellow). Bottom panels show high-magnification view of top panels. Scale bars represent 20 μm unless otherwise noted.

Sterol O-Acyltransferase 1 (Soat1) esterifies excess intracellular FC into CE for storage in LDs ^44^. We therefore generated an overexpression line for the *Drosophila soat1* ortholog, *CG8112*. Co-expression of Soat1 with Fru significantly reduced FC accumulation and fibrosis-like phenotype while restoring LD abundance (Fig. 5E). Together, these findings indicate that JAK/STAT and its downstream effectors Fru and Hsl mediated LD remodeling is essential for maintaining cholesterol homeostasis and preventing lipotoxic stress in steroidogenic cells.

### Peripheral leptin-family signals orchestrate lipid remodeling to drive sexual maturation

To investigate the source of the systemic signal and identify developmental cues that regulate JAK/STAT activation in the PG, we examined tissue-specific expression of Upd2 and Upd3, the fly leptin analogs and ligands of the JAK/STAT signaling pathway that mediate long-range endocrine signaling ^45, 46^. Immunostaining with an anti-Upd3 antibody revealed broad expression across larval tissues, including the fat body, muscles, brain, imaginal discs, and intestine, with levels progressively declining from mid- to late-L3 (Fig. 6A). Consistent with these protein dynamics, RT-qPCR confirmed that *upd2* and *upd3* transcripts in imaginal discs decreased from mid- to late-L3 (Fig. 6B).

**Figure 6.**
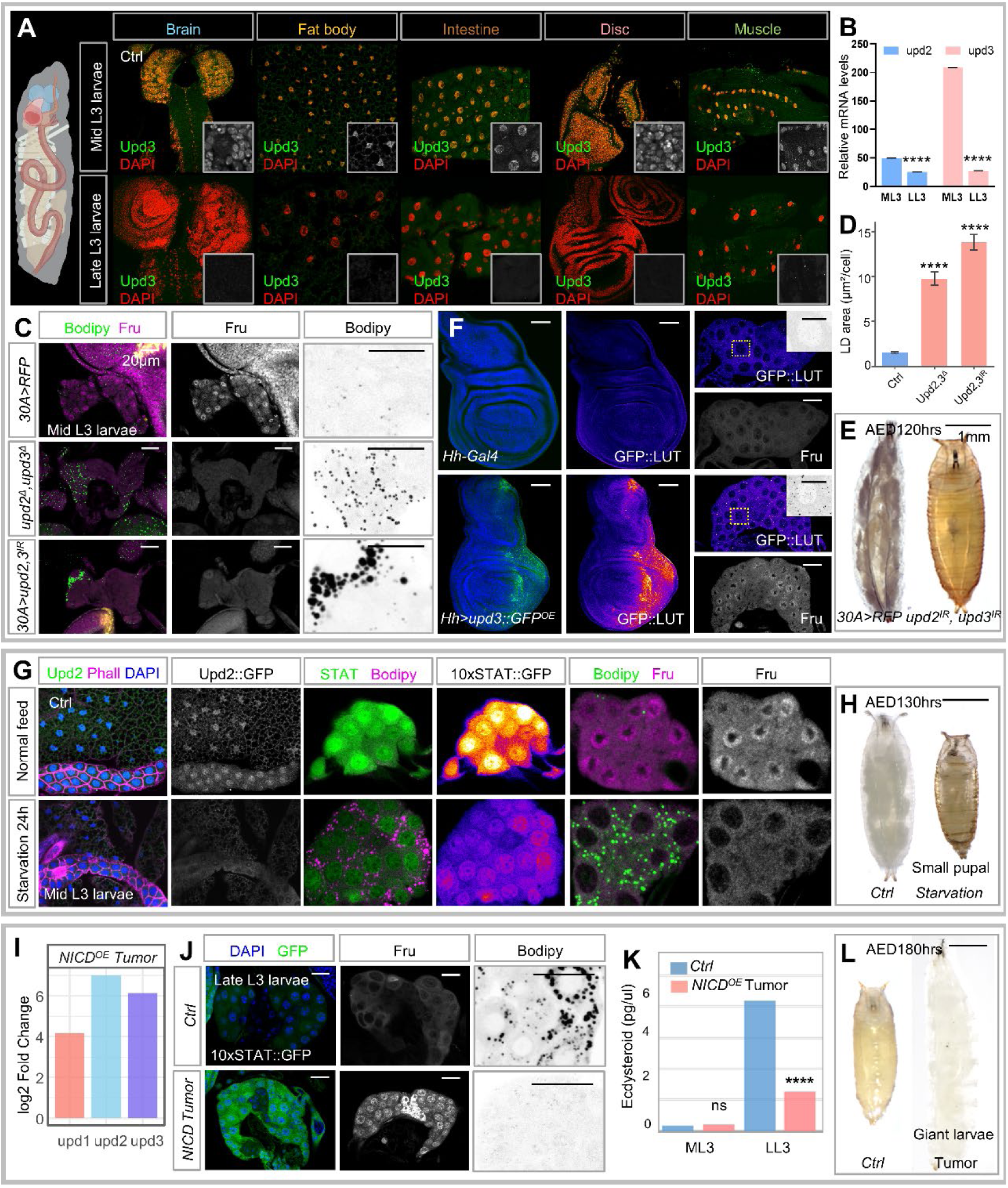
Inter-organ Upd signaling orchestrates the JAK/STAT-Fru-Hsl axis in PG cells. (A) Confocal images of various tissues including brain, fat body, intestine, wing disc, and muscle from mid-L3 and late-L3 larvae. Upd3 (green) is widely expressed in mid-L3 but significantly downregulated by late-L3. DAPI (red) marks nuclei. (B) qPCR analysis showing the relative mRNA levels of *upd2* and *upd3* at mid-L3 and late-L3. (C) Confocal images of Fru (magenta) and LD (Bodipy, green) staining in PG cells at mid-L3 larvae under imaginal disc specific (*30A-Gal4*) knockdown of *upd2*, *upd3*, or simultaneous depletion of *upd2* and *upd3*. (D) The graph shows quantification of LD area in the PG based on images (C) normalized to controls. Representative larvae and prepupal images (E) at 120 h after egg laying, including control (*30A>RFP*), imaginal disc-specific knockdown of *upd2*, *upd3*. (F) Representative images showing GFP distribution upon *Hh-Gal4* driven Upd3-GFP overexpression in the wing disc (left) and the PG (right). Grayscale images showing Fru expression level. (G) PGs from normally fed and 24h-starved larvae. Starvation leads to a marked decrease in Upd2 levels, reduced STAT activity (10xSTAT::GFP), loss of Fru expression (magenta), and massive accumulation of LDs (green). (H) Morphological comparison showing small pupae resulting from starvation at 130 hrs AED. (I) RNA-seq data showing upregulated expression of *upd1*, *upd2*, and *upd3* in NICD overexpression tumor. (J) PGs from larvae bearing NICD tumors show constitutive STAT activation (10xSTAT::GFP), and elevated Fru expression (gray), and premature depletion of LDs (Bodipy). (K) Ecdysteroid titers (pg/µl) determined by ELISA in control (*Hh-Gal4*) and *NICD^OE^* tumor-bearing larvae at mid-L3 and late-L3. (L) *NICD* tumor-bearing larvae exhibit a “giant larvae” phenotype and failure to pupariate at 180hrs AED. Data are presented as mean ± s.e.m. Statistical significance was determined by one-way ANOVA with multiple comparisons. ****p < 0.0001; ns, not significant. Scale bars represent 20μm (PG), 50μm (wing disc), and 1 mm (larvae).

To test whether systemic Upd2/Upd3 signaling modulates PG metabolism, we analyzed LD levels in an *upd2^Δ^, upd3^Δ^* double mutant and observed increased LD accumulation in PG cells at mid-L3 relative to controls (Fig. 6C, D). Similarly, imaginal disc-specific knockdown of *upd2* and *upd3* (driven by *30A-Gal4*) phenocopied LD accumulation in the PG (Fig. 6C, D). These effects were accompanied by reduced Fru expression at mid-L3, indicating that Upd-dependent LD remodeling is mediated by the JAK/STAT-Fru-Hsl axis (Fig. 6C). Functionally, these animals exhibited precocious developmental progression, entering pupation earlier than controls at 120 h AED (Fig. 6E).

Consistently, ectopic expression of *upd3-GFP* in imaginal disc using multiple Gal4 drivers (*Hh-Gal4*, *Ci-Gal4*, and *pdm2-Gal4*) supported non-cell-autonomous signaling, as GFP signal and Fru upregulation were both detected in PG cells (Fig. 6F and Fig. S7A-C). Together, these data indicate that imaginal tissues secrete dynamically regulated Upd/leptin-family signals that systemically tune steroidogenic lipid metabolism, thereby coupling inter-organ growth cues to metabolic control of sexual maturation.

### Systemic leptin-family signals integrate nutritional and pathological stress to tune steroidogenic output

In mammals, circulating leptin levels closely track nutritional status, decreasing during nutrient deficiency and increasing under nutrient sufficiency ^47, 48^, and Upd signaling in adult *Drosophila* shows conserved nutrient responsiveness ^45^. To determine whether the larval PG engages the JAK/STAT-Fru-Hsl brake to sense nutritional state, we subjected larvae to starvation during mid-L3. Nutrient deprivation markedly reduced Upd2 expression level in both the fat body and salivary gland (Fig. 6G). Correspondingly, JAK/STAT activity in the PG declined, promoting nuclear Fru loss and premature release of LD storage constraints (Fig. 6G). Functionally, starved animals entered pupation within 24 hours and produced smaller pupae relative to fed controls (Fig. 6H). These findings indicate that the PG detects systemic peptide hormone fluctuations under nutrient stress and accelerates steroidogenesis as an adaptive response.

Beyond nutritional cues, inflammation is a potent inducer of leptin/Upd signaling, particularly during malignancy and infection ^49–52^. To determine whether pathological states perturb steroid homeostasis through the Upd-JAK/STAT-Fru-Hsl axis, we performed RNA-seq analysis on a *Drosophila* tumor model induced by NICD overexpression in the salivary gland imaginal ring (ImR) ^53, 54^, which revealed significant upregulation of *upd1*, *upd2*, and *upd3* (Fig. 6I). Consistent with this, JAK/STAT activity was strongly elevated in PGs of tumor-bearing larvae, accompanied by an increased Fru expression and enhanced LD hydrolysis (Fig. 6J). Functionally, tumors disrupted systemic steroidogenesis: ELISA showed that *NICD^OE^* tumors blunted the normal late-L3 rise in ecdysteroid titers, thereby blocking the larval-to-pupal transition (Fig. 6K). *NICD^OE^* tumor-bearing larvae failed to pupate and persisted as giant larvae for up to ∼280 h AED, compared with controls that pupated at ∼130 h AED (Fig. 6L). Notably, moderate starvation partially rescued developmental arrest in *NICD^OE^*tumor-bearing larvae, allowing LDs to reaccumulate in the PG after 36 hours of starvation and enabling giant larvae to initiate pupariation (Fig. S8A, B). To evaluate the generalizability of these findings, we extended our transcriptomic profiling to three additional *Drosophila* tumor models, including *lethal giant larvae* (*lgl*), *scribble* (*scrib*) or *Snf5-related 1* (*snr1*) knockdown in wing discs ^55, 56^. RNA-seq analysis across all three models revealed a consistent upregulation of the *upd* family ligands, which subsequently triggered elevated Fru expression in PG cells (Fig. S8C, D).

Taken together, these findings establish Upd ligands as peripheral tropic signals that directly modulate the JAK/STAT-Fru-Hsl axis within steroidogenic cells. By linking developing tissues and tumor-derived cytokines to PG metabolism and steroid hormone output, the Upd/leptin family integrates nutritional state, tissue stress, and developmental growth signals to precisely control sexual maturation timing.

### Cross-species analyses reveal conserved features of steroidogenic metabolic reprogramming

Given the conserved LD dynamics across steroidogenic tissues, we next examined whether the upstream regulatory program is likewise conserved. To test whether the JAK/STAT-Fru-Hsl module is conserved in the PG of the hemimetabolous insect *Blattella germanica*, we performed RNA-seq profiling of PG tissue from early- and late-L6 nymphs. This analysis revealed broad transcriptional remodeling of lipid metabolic programs. Notably, key components of the JAK/STAT-Fru-Hsl axis, including *Socs36E*, *dome*, and *fru*, as well as lipases (*Lip3*, *Lip4*), were developmentally downregulated. In contrast, genes associated with ecdysteroid biosynthesis, including *dib*, *nvd*, *ftz-f1*, and *Eip93F*, were coordinately upregulated, consistent with progression through the developmental transition (Fig. 7A). These findings suggest that JAK/STAT-Fru-Hsl-mediated lipid remodeling in the PG is potentially conserved across both holometabolous and hemimetabolous insects.

**Figure 7.**
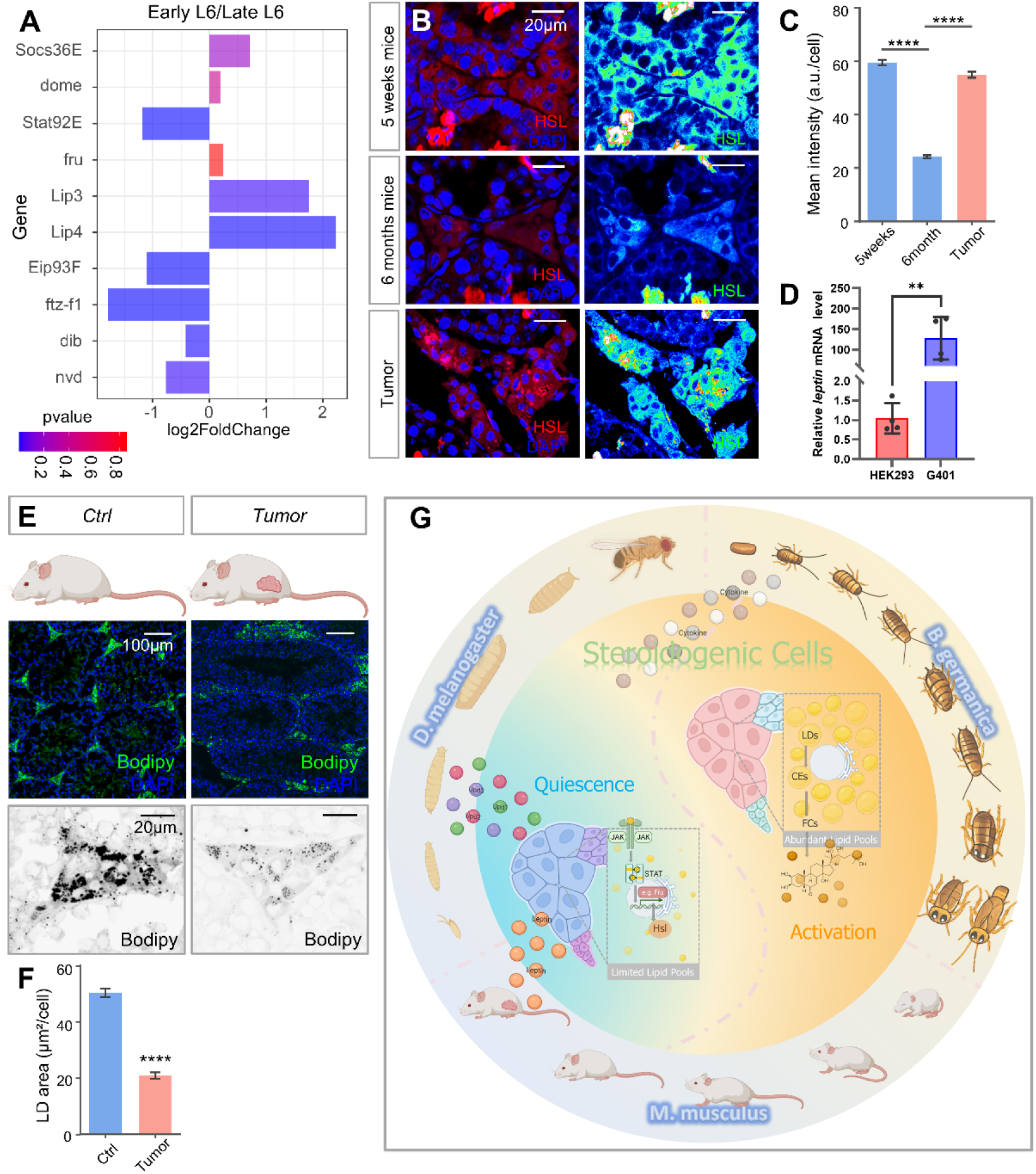
Conservation of the leptin-JAK/STAT-HSL axis in both insects and mammalian steroidogenesis. (A) Fold-change plot of differentially expressed genes (DEGs) is shown with log2 fold change values and p-value color coding from RNA-seq analysis comparing early versus late L6 nymph in the PG. (B) Representative confocal images and quantification (C) of HSL protein levels in mouse Leydig cells from young (5 weeks), adult (6 months), and tumor-bearing mice. Scale bar: 20 μm. (D) qPCR analysis of relative *leptin* mRNA levels in HEK293T cells compared to G401 cells, showing significant upregulation of leptin signaling in the tumor context. (E) Bodipy staining (green) of steroidogenic tissues in control (6 months) and tumor-bearing mice (4 months). (F) Quantitative analysis of LD area in Leydig cells based on image E. (G) A schematic model illustrating the mechanism across *D. melanogaster*, *B. germanica*, and *M. musculus*. The leptin/Upd-JAK/STAT-HSL axis acts as a universal spatiotemporal switch, coordinating lipid mobilization to transition steroidogenic cells from a quiescent (LD-poor) to an activated (LD-rich) state during development or tumor-induced stress. Data are presented as mean ± s.e.m.. Statistical significance was determined by one-way ANOVA followed by multiple comparisons. **p < 0.01, ****p < 0.0001.

To further examine the evolutionary conservation of this regulatory logic, we analyzed HSL expression in developing mouse testis. During normal maturation, immunofluorescence analysis showed that HSL expression in Leydig cells declined with age, with signal intensity at 6 months significantly lower than at 5 weeks (Fig. 7B, C), paralleling developmental changes observed in *Drosophila* (Fig. 4E, F). Consistent with this trend, LD size and number were developmentally regulated in mammalian steroidogenic tissue (Fig. 1A, B).

We next asked whether this conserved program is engaged under pathological conditions. Because loss of the tumor suppressor *Snr1* (the *Drosophila* ortholog of SMARCB1) activates the Upd-JAK/STAT-Hsl regulatory axis (Fig. S8C, D), we investigated whether *SMARCB1*-deficient tumors ^57^, which underlies pediatric atypical teratoid/rhabdoid tumors (ATRT) and malignant rhabdoid tumors (MRT), similarly engage leptin-associated systemic signaling. Analysis of the SMARCB1-null rhabdoid tumor cell line G401 revealed an approximately 130-fold increase in leptin expression relative to a control cell line (HEK293) (Fig. 7D). To extend these findings in vivo, we established a mouse xenograft ATRT model by subcutaneously injecting G401 cells into immunodeficient mice and asked whether pathological stress perturbs this program. Tumor-bearing mice exhibited a marked reduction in LD size and number in Lydig cells (Fig. 7E, F), suggesting that systemic effects of cancer disrupt LD storage in mammalian steroidogenic tissues, a phenotype reminiscent of that observed in tumor-bearing *Drosophila*. Consistently, immunofluorescence analysis showed that HSL levels were significantly elevated in Leydig cells of tumor-bearing mice (Fig. 7B, C), indicating enhanced lipolytic activity. Notably, although mammals lack a Fru ortholog ^58^, these findings support partial conservation of leptin-responsive JAK/STAT-HSL-associated lipid remodeling in steroidogenic cells.

In summary, our study identifies a partially conserved feedback circuit in which leptin-family signals from peripheral tissues are integrated within steroidogenic organs to drive rapid JAK/STAT-lipase-mediated reprogramming of LD-CE homeostasis. This mechanism fine-tunes steroid hormone output and aligns sexual maturation with physiological state. More broadly, our findings provide a framework for understanding how inter-organ endocrine communication translates systemic growth and stress cues into metabolic decisions within steroidogenic tissues, thereby promoting developmental robustness while preserving adaptability to pathological and environmental change (Fig. 7G).

## Discussion

The coordination of sexual maturation with developmental transitions requires endocrine systems to precisely integrate systemic growth cues with steroid hormone production. In this study, we define in *Drosophila* a leptin/Upd-JAK/STAT-Fru-Hsl signaling axis that functions as a metabolic gatekeeper within steroidogenic tissues to regulate the timing of metamorphosis. Key components, particularly LD-dependent cholesterol mobilization and Hsl-mediated CE hydrolysis, are highly conserved, suggesting broader physiological relevance.

Steroidogenic tissues possess limited preformed hormone stores and therefore rely on the dynamic mobilization of stored CEs from LDs to sustain the rapid production of steroid precursors required for developmental transitions. We show that a developmental decline of the systemic cytokine Upd, a functional analog of mammalian leptin, drives a stage-specific reduction in JAK/STAT signaling within the PG, the insect steroidogenic organ. Elevated JAK/STAT activity during early-L3 larval growth acts as a molecular brake that restricts LD accumulation, thereby limiting cholesterol precursor availability and preventing precocious maturation. As systemic Upd/leptin levels decrease, this inhibitory constraint is released, permiting the accumulation of cholesterol-rich LDs. This metabolic shift provides the substrate necessary to initiate ecdysone biosynthesis and trigger the transition to adulthood. Together, our findings establish a paradigm in which inter-organ peptide hormone circuit governs the upstream metabolic state of a steroidogenic organ, revealing an unrecognized checkpoint in developmental timing.

This mechanism complements classical tropic hormone pathways in animals. In mammals, puberty and stress responses are primarily driven by pituitary trophic factors (LH/FSH for gonads, ACTH for adrenals), which act through well-characterized second-messenger cascades ^14, 59^. In rodents, most of the cholesterol required for steroidogenesis in the adrenal gland, ovary, and testicular Leydig cells is acquired via SR-BI mediated selective uptake of HDL-cholesterol ^60, 61^, whereas in humans the majority of cholesterol for steroid synthesis is supplied through the LDL-receptor-dependent endocytic pathway ^14, 62^. These mechanisms primarily mobilize cholesterol from extracellular lipoproteins or through *de novo* synthesis. In contrast, the leptin/Upd-JAK/STAT-HSL axis mobilizes intracellular cholesteryl esters stored in LDs. Functionally, leptin/Upd acts as a peripheral tropic signal that modulates steroid outputs, complementing canonical neuroendocrine pathways (HPG/HPA in vertebrates and PTTH in insect) by enabling a more rapid metabolic-endocrine response to developmental and stress cues. Notably, although mammalian leptin and related cytokines are known to modulate the HPG axis ^63^, our findings uncover a direct cytokine-mediated link to steroidogenic cells, bypassing the multi-tiered neuroendocrine cascade.

In this metabolic-endocrine coupling pathway, the rapid remodeling of LD-CE stores in steroidogenic cells is a key response to inter-organ developmental and nutritional signals. In all steroidogenic cells, the LD core is enriched in CEs, which forms a reservoir of hormone precursors ^9^. Mobilization of FC from these stores is a rate-limiting step in hormone synthesis: in vertebrates, HSL has long been recognized as the major nonspecific CE hydrolase, supplying FC for mitochondrial conversion ^9^. Indeed, HSL-mediated LD-CE hydrolysis is known to deliver cholesterol to StAR in the mitochondria of adrenal and gonadal cells ^64^. Given that HSL can also act as a lipase to cleave fatty acids from DG and thereby facilitate LD lipolysis, we found that leptin/Upd-JAK/STAT signaling upregulates HSL expression in steroidogenic cells outside developmental stages, accelerating LD catabolism and suppressing LD storage. This fine-tuned metabolic decision ensures efficient allocation of limited resources. Consistently, HSL loss led to LD accumulation (Fig. 4G) and impaired steryl ester mobilization in flies ^65^, as well as LD buildup in the mouse adrenal cortex accompanied by markedly reduced corticosterone levels ^66^, male infertility, and reduced testis weight ^67^, indicating that HSL is an evolutionarily conserved regulator of steryl ester turnover. Thus, LD-CE remodeling emerges as a critical rate-limiting step in developmental steroid output.

Complementing HSL function, the cholesterol esterifying enzyme SOAT1 acts antagonistically to maintain proper cholesterol turnover and timely availability within steroidogenic tissues. SOAT1 is evolutionarily highly conserved, being present from yeast to humans ^68^, and is robustly expressed in steroidogenic tissues, including Leydig cells, ovarian granulosa cells, and the adrenal cortex ^69^. In animals, SOAT1 overexpression promotes the accumulation of LDs and CEs ^70^, whereas its inhibition enhances FCs availability and subsequent testosterone synthesis ^71^. Our study demonstrates that Soat1 overexpression effectively mitigates the pathological FC accumulation and LD depletion caused by hyperactivation of the leptin/Upd-JAK/STAT-Hsl axis by converting excess intracellular FC into CEs for storage within LDs. These findings indicate that Soat1 and Hsl constitute a fundamental homeostatic rheostat orchestrating the precise CE-FC turnover required to modulate circulating steroid hormone levels.

Although high levels of leptin and JAK/STAT signaling have been reported to suppress steroid hormone production ^72–75^, the precise molecular mechanisms remain unclear, and our findings suggest that the leptin/Upd-JAK/STAT-Hsl axis functions as a central hub integrating diverse physiological and pathological conditions, demonstrating high adaptability and flexibility. Starvation, which lowers systemic leptin/Upd levels, prematurely releases this molecular brake, leading to accelerated maturation as an adaptive strategy to ensure reproductive success under adverse nutritional conditions. Notably, the surge in steroid hormone production accompanying sexual maturation is also initiated when larvae cease feeding, and the mechanism uncovered here likely governs this physiological transition as well. In contrast, pathological conditions such as tissue damage and tumorigenesis induce hypersecretion of Upd/leptin cytokines ^38, 76^ (Fig. 6I). Sustained, nonphysiological activation of the JAK/STAT-Fru-Hsl axis results in chronic LD depletion and suppressed ecdysteroid titers, leading to severe developmental arrest and a giant larva phenotype (Fig. 6J-L). Consistently, we observed chronic LD depletion and high HSL expression in the gonads of tumor-bearing mice (Fig. 7B-F). This mechanism provides crucial molecular insight into how inflammatory and cancer-related cytokines, a well-established component of systemic inflammation and cancer cachexia in mammals, can disrupt the hormonal control of development and growth. Therefore, targeting this conserved axis presents a promising avenue for therapeutic intervention, both for disorders of developmental timing (e.g., precocious or delayed puberty) and for mitigating endocrine dysfunction associated with systemic disease.

## Materials and Methods

### Fly strains and genetics

Flies were maintained at 25 °C, 60% relative humidity, and 12-h light/dark cycle. Flies were reared on Bloomington *Drosophila* Stock Center (BDSC) cornmeal and yeast-based diet, unless otherwise noted. The Gal4/UAS driven RNAi crosses were cultured at 25°C until eclosion, and the same culture conditions were also used for the control group. Tub-Gal80ts was used to control Gal4 activity. *UAS-fru^A^* and *UAS-fru^B^* were gift from Dr. M. Arbeitman. Detailed genotypes were listed in table S1.

### RNA-seq and data analysis

Larvae ring glands were carefully dissected in 1 × PBS before RNA extraction. Total RNAs were extracted from L3 stage ring glands using the Zymo RNA preparation kit. Biological replicates were collected in triplicate for tumor samples and in duplicate for ring gland samples per genotype. NEBNext Poly(A) mRNA Magnetic Isolation Module and NEBNext Ultra II RNA Library Prep Kit for Illumina were used for library preparation. The libraries were sequenced using an Illumina HiSeq 2500 system, obtaining 40 million reads for each sample.

Raw reads were preprocessed using fastp (v0.22.0) with default parameters to ensure high-quality data retention ^77^. This resulted in the retention of a vast majority of the bases and reads, with minimal differences among all libraries. Subsequently, we employed bowtie2 - an intron-aware short-read aligner - to map all reads to the dme6 reference genome ^78^. To quantify gene expression levels, we utilized RSEM 1.2.31 and performed pairwise group comparisons using DESeq2 1.44.0 on the output generated by RSEM ^79, 80^. Significantly differentially expressed genes (DEGs) were selected based on the threshold of p-value ≤ 0.05 and ≥2-fold change. The alignment of reads and generation of the differentially expressed gene (DEG) matrix were carried out using the scripts provided by Trinity RNA-Seq ^81^.

To align the *Blattella germanica* transcripts to those of *Drosophila*, we performed de novo transcriptome assembly using the Trinity RNA-Seq v2.14.0 pipeline ^82^. After assembly, the transcripts were aligned to the *Drosophila* transcriptome (GCF_000001215.4) using BLAST+ v2.12.0+. The assembled transcripts were then used as a reference library for gene expression quantification and differential expression analysis, following the standard Trinity pipeline.

After generating the expression matrix, we performed sample clustering using principal component analysis (PCA) from the R package ^83^, and utilized the “pheatmap” function to generate heatmaps. Downstream functional insights were gained via (GSEA) ^84^, using clusterProfiler 4.2 ^85^. The KEGG pathway maps were visualized using pathview^86^.

### cDNA Synthesis and RT-qPCR

Larvae tissues were dissected in 1 × PBS and stored at −80°C until all timepoints were collected. Isolated tumors and control cells were directly preserved in TRIzol reagent (Thermo Fisher Scientific). The samples were homogenized with TRIzol (Thermo Fisher Scientific) using standard protocols on ice. Following phase separation, the aqueous phase was transferred into a new tube, mixed with an equal volume of 70% ethanol and loaded directly onto the mini kit columns (Zymo RNA preparation kit). The remaining steps of the RNA isolation were performed in accordance with the manufacturer’s protocol, including on-column DNase digestion (Qiagen) for 15 minutes at room temperature.

Reverse transcription was performed on 0.25-1μg total RNA using the Superscript Reverse Transcriptase II from Thermo Fisher Scientific (Cat# 18064022) and oligo(dT) primers. cDNA was used as a template for qPCR.

RT-qPCR was performed with a Quantstudio-3 Real-Time PCR System and PowerUp SYBR Green Master Mix (Thermo Fisher Scientific). Three independent biological replicates were performed with three technical replicates each. The mRNA abundance of each candidate gene was normalized to the expression of GAPDH by the comparative CT methods. Primer sequences are listed in Table.S2.

### Generation of UAS-SOAT Transgenic Flies

To generate transgenic flies for UAS-SOAT, we amplified the open reading frame corresponding to the *CG8112-RB* isoform using the cDNA clone RE61294 (*Drosophila* Genome Resource Center; DGRC, Bloomington, IN) as template and the primers 5’-TAATGAATTCATGGCCGCTGAAAGGCAGG-3’ and 5’-TAATGCGGCCGCCTAATTGTA GCAGCGCCA-3’. The parameters of the PCR were 5 min 98 °C; 35 cycles of 30 sec 98 °C, 30 sec 55 °C, 90 sec 72 °C; 10 min 72 °C; hold at 4 °C. An aliquot of the PCR product was then resolved on a 0.8% agarose gel supplemented with ethidium bromide at 90 V. The PCR product and the plasmid pCaSpeR3 were enzymatically digested by the restriction enzymes EcoRI and NotI in CutSmart Buffer for 2 h at 37 °C followed by heat inactivation of the enzymes at 65 °C for 30 min. The digested vector DNA was resolved on a 0.8% agarose gel at 90 V for 45 min and isolated using the Wizard SV Gel Clean-Up System (Cat#: A9281, Promega, Madison, WI) according to the manufacturer’s instructions. Subsequently, the restriction products were ligated for 2 h by 37 °C in 10 µl T4 DNA-Ligase Buffer containing 0.5 µl T4 DNA Ligase (NEB). E. coli XL-1 were electroporated with 3 µl of the ligated plasmid and plated on LB plates with 100 µg/ml Ampicillin. Positive colonies were identified by plasmid miniprep and sequencing, amplified and isolated by Midiprep using the NucleoBond Xtra Midi kit (Cat#: 740410.50, Macherey-Nagel, Düren, DE) according to the manufacturer’s instructions. This plasmid preparation was injected into a *w^1118^* host strain and transgenic flies were established by Bestgene (Chino Hills, CA). Homozygous stocks of the genotypes w1118; *P{w+mC=UAS-SOAT} M2* and w1118; +; *P{w+mC=UAS-SOAT}M5* with transgene insertions on the 2^nd^ and 3^rd^ chromosome, respectively, were subsequently established.

### Immunostaining and Confocal Imaging

Tissue samples were dissected in phosphate buffered saline (PBS), then fixed in 4% formaldehyde in PBS for 30 minutes. After washing with PBS with 0.1% Triton X-100 (PBT), the samples were incubated in PBT with primary antibodies at 4°C overnight with shaking and then washed in PBT three times for 15 minutes each. The following antibodies were used in immunostaining: anti-GFP (Cell Signaling Technology, #2956S, 1:200), anti-β-gal (Promega, #PAZ3783, 1:500), anti-Hsl (Cell Signaling Technology, #4107S, 1:200), anti-Fru^COM^ rabbit polyclonal antibody (1:500) ^34^, and anti-Upd3 rat monoclonal antibody (1:50) ^87^. The secondary antibodies conjugated with Alexa 488, Alexa 546 or 633 (Invitrogen) were diluted 1:200 and incubated at room temperature for 2 hours.

Fixed tissues were co-stained with wheat germ agglutinin conjugated to Alexa Fluor 555 (WGA-Alexa, 1 μg/ml, Molecular Probes, Eugene, OR) to visualize fibrotic structures. Nuclei were labeled with DAPI (Invitrogen, D1306, 1:1000). After washing, samples were mounted and imaged with Zeiss LSM 900 or Zeiss LSM 980 Confocal Microscopes. Mean fluorescence intensity from each section was quantified using ImageJ (Fiji) software ^88^.

### Filipin Cholesterol Staining

For in-situ filipin staining, staged ring glands from control and overexpression groups were dissected and fixed for 15 min in 4% paraformaldehyde, followed by washes in PBS (three times for 15 min). A fresh stock solution of filipin (1 mg/ml in DMSO; Sigma-Aldrich) was diluted to a final concentration of 50 μg/ml in PBS. Tissues were incubated in the dark for 30 min at room temperature and rinsed with PBS three times.

Tissues were mounted in Vectashield (Vector Laboratories) and immediately visualized with Zeiss LSM 900 confocal scanning microscope equipped with a UV laser.

### Alizarin Red S Staining

Tissues were fixed in 10% neutral buffered formalin for 30 min, followed by two rinses with distilled water. Samples were then transferred onto clean glass slides and incubated with 2% Alizarin Red S solution (pH 4.3, Merck Millipore, USA) for 30 min at room temperature. After three subsequent washes in PBT, the ring glands were mounted in Elvanol. Bright-field imaging was performed using a Zeiss LSM 980 microscope under brightfield illumination (and a 592 nm bandpass filter).

### Starvation

Larvae were collected within 2 hours after entering the L3 stage and rinsed with water to remove residual food. For each genotype, larvae were randomly assigned to either fed or starved groups. Fed groups (20 larvae per vial, 3 vials) were maintained on standard fly food, while starved groups were transferred to vials containing 1% non-nutritive agar. Following a 24hr starvation period at 25°C, larvae were dissected and processed for staining alongside fed controls. For tumor-bearing models, starvation was initiated 4 days after tumor induction, with both experimental and control groups maintained under the same conditions at 29 °C.

### Lipid Stains

Ring glands dissected from larvae were fixed in 4% paraformaldehyde for 30 min at room temperature, rinsed in PBS, and incubated in Bodipy 493/503 (1:1000, Thermo Fisher Scientific Cat#D3922) or NileRed (1:2000, TCI America, 7385-67-3) for 1hr at room temperature, in the dark, to stain the neutral LDs. Next, the ring glands tissues were rinsed in PBS and mounted in antifade reagent with DAPI.

### Time-lapse Imaging

Larvae were first dissected carefully in M3 culture medium to isolate the brain-ring gland complexes. The complexes were then transferred to a glass-bottom culture dish (MatTek) with a drop of M3 medium, after which the brain tissues were quickly separated from the ring glands and removed from the dish. A coverslip was then positioned on top of the ring glands to prevent sample movements, and more M3 medium was added to the dish before imaging. The samples were imaged with a Zeiss LSM 980 confocal microscope using the time-lapse configuration.

### Transmission electron microscopy

Larvae were dissected in PBS buffer and fixed in fixative containing 2% paraformaldehyde and 2.5% glutaraldehyde in cacodylate buffer. Then samples were post-fixed in 1% osmium tetroxide for 1 hour, dehydrated progressively in acetone, and embedded in Poly/Bed 812 epoxy resin. Following heat polymerization, resin blocks were sectioned using Leica UC7 microtome toward 70 nm ultra-thin sections. Finally, the section grids were post-stained by uranyl acetate and lead citrate and then imaged using FEI Tecnai G2 F30 transmission electron microscope.

### Rescue experiments with 20E

5 mg/mL stock solution of 20-hydroxyecdysone (20E) (Sigma-Aldrich) was prepared by dissolving 5 mg of powder in 1 mL of EtOH. Working solution was adjusted by diluting the initial stock solution in PBS until a final concentration of 0.3 mg/ml. Control solutions were prepared using the same amount of EtOH in PBS. The total volume for both 20E and control solutions was adjusted to 100 mL. Adult flies were placed on either standard fly food or food supplemented with active 20E. Following 10h egg-laying period at 25°C, adult flies were removed, and the development of the larvae was monitored.

### JAK/STAT inhibitor

The JAK2 inhibitor of AG490 (T-9142, LC Laboratories) was dissolved into DMSO (25 mg/ml) and 100 µl was added directly on the surface of fly vial food to reach a final concentration of 250 ng/ml. Larvae were collected within 2 hours after entering the L3 stage and rinsed with water to remove residual food. Larvae were then transferred in groups of 20 to vials containing either standard food or AG490-supplemented food.

Following 24h incubation at 25°C, ring glands were harvested for further analysis.

### Fluorescence analysis of cholesterol distribution by 22-NBD-cholesterol

To assess the incorporation of cholesterol and to visualise its distribution, we conducted *in vitro* incubation of the brain-ring gland complexes dissected from third instar larvae with 22-(*N*-(7-Nitrobenz-2-Oxa-1,3-Diazol-4-yl) Amino)-23,24-bisnor-5-cholen-3β-ol (22-NBD-cholesterol; Life Technologies). 22-NBD-cholesterol was dissolved in 100% ethanol at a 2 mM concentration for a stock solution. The third instar larvae were dissected in PBS and the brain-ring gland complexes were transferred into Schneider’s *Drosophila* Medium (Life Technologies) containing 10% fatal bovine serum, 100 U/ml penicillin (Wako) and 100 µg/ml Streptomycin (Wako). After an incubation at 25°C for 10 min, the medium was replaced with a fresh medium containing 0.5% 22-NBD-cholesterol stock solution, which achieved a 10 µM 22-NBD-cholesterol with 0.5% final ethanol concentration. Then, tissues were incubated at 25°C for 5 hours in a dark condition. Tissues were washed with PBS twice and mounted. A 488 nm laser was used for excitation of 22-NBD-cholesterol fluorescence and fluorescence emission was selected by a 490–555 nm-band pass filter. Fluorescence images were obtained with an LSM 980 laser-scanning confocal microscope (Zeiss).

### Ecdysteroid titer measurement (ELISA)

The ecdysteroid titers of larvae were measured using the 20-hydroxyecdysone Enzyme Immunoassay (EIA) kit, (Cayman Chemicals) which detects both ecdysone and 20-hydroxyecdysone. Briefly, frozen larvae were homogenized in methanol and ecdysteroids were extracted as described previously ^89^. The extracts were evaporated in a Speed Vac and the residue resuspended in EIA buffer (Cayman Chemical) and analyzed following the manufacturer’s protocol. A standard curve was determined using a dilution series containing a known amount of purified 20-hydroxyecdysone solution provided by the kit. Absorbance at 415 nm was detected using a benchtop microplate reader (Bio-Rad).

### Mammalian cell culture

HEK293 (ATCC® CRL-1573TM, ATCC) and G401 (ATCC, CRL-1441TM, ATCC) cell lines were obtained from Dr. Sang’s lab at FSU ^90^.

HEK293 cells were cultured in DMEM supplemented with 10% fetal bovine serum (FBS) at 37 °C with 5% CO₂. G401 cells were cultured in McCoy’s 5A medium (ATCC, 30-2007™) supplemented with 10% FBS.

### Xenograft mouse model

All mouse studies were conducted in accordance with the guidelines and regulations approved by the Institutional Animal Care and Use Committee at Tulane School of Medicine. Mice received standard care and were euthanized according to established Institutional Animal Care and Use Committee protocols. Xenograft experiments were performed in 3-month-old immunocompromised nude male mice (The Jackson Laboratory). To induce tumors, G401 cells (1,000,000 cells) were transplanted into the flank of each mouse. Mouse health was monitored every two days. Four weeks later, samples were harvested.

### Mouse testis cryosectioning

Testis were fixed in freshly prepared 4% paraformaldehyde overnight, washed three times with 1× PBS, and then stored in 30% sucrose at 4°C overnight. The testes were oriented in Tissue-Tek (Sakura Finetek, Japan), embedded in OCT, and sectioned at 10 μm using a cryostat ^91^. Mouse testis sections were washed three times with PBS, and stained with 1 μg/mL Bodipy 493/503 (Invitrogen, D3922) for 2 hrs. After three rinses in PBS, the sections were mounted in antifade mounting medium with DAPI.

### Immunohistochemistry for the mouse testis

Antigen retrieval was performed by steaming the slides in 10 mM sodium citrate. After cooling, endogenous peroxides were inactivated by incubating the slides in 3% hydrogen peroxide for 10 min. Tissues were permeabilized in 0.1% PBS-Triton X-100 (PBST) for 30 min with low-speed shaking, blocked with 5% NGS in 0.02% PBST at room temperature for 1 h, and incubated with primary antibodies diluted in 2.5% NGS with 0.02% PBST at 4°C overnight. After 3 times of 0.02% PBST wash, tissues were incubated with the secondary antibodies diluted in 2.5% NGS (in 0.02% PBST) at room temperature for 1 h, followed by 3 times of 0.02% PBST wash. The stained tissues were stored in PBS at 4°C until imaging. During the staining process, the tissues were shaken gently to preserve the lipid signal.

### Quantification and Statistical Analysis

Data analyses were conducted in GraphPad Prism and the R package. Unpaired t-test was used for two-sample comparisons, one-way ANOVA Dunnett’s multiple comparison test and One-way ANOVA with Tukey’s multiple comparison test were used for multiple-sample comparisons. Specific statistical approaches for each figure are indicated in the figure legend. Non-normally distributed data, foraging index for instance, were analyzed using the Wilcoxon rank-sum test to determine the significant difference between groups. The statistical test used for each figure is clarified in the figure legend. The raw mean, SEM and p-value are presented in supplementary tables.

## Supporting information

Supplemental vedio 1

Supplemental vedio 2

Supplemental vedio 3

Supplemental vedio 4

## Acknowledgements

We thank Dr. M. Arbeitman, Bloomington *Drosophila* Stock Center (BDSC, USA) and the Vienna *Drosophila* RNAi Center (VDRC, Austria) for providing fly stocks; Dr. R. Niwa, MB. O’Connor, S. Ohsawa, Y. Yagi and the Developmental Studies Hybridoma Bank (DSHB, USA) for providing antibodies. We also thank QY. Zhang for providing mouse samples. Special thanks to A. Brown from the FSU Department of Biological Science for help with bulk RNA-seq library preparation.

## Authors’ contribution

Conception and design: JS, WMD

Development of methodology: JS, WMD

Acquisition of data: JS, CE, AA, CH, XFW, FLH, YCH

Analysis and interpretation of data: JS, WKL, HCB

Study supervision: JS, WMD

Writing – original draft: JS, WMD

Writing – review & editing: JS, WMD

Acquisition of funding: WMD

All authors read and approved of the final manuscript.

## Declaration of Generative AI and AI-assisted technologies in the writing process

During the preparation of this work the authors used ChatGPT to check for grammar and English usage. After using this tool/service, the author(s) reviewed and edited the content as needed and take full responsibility for the content of the publication.

## Data and materials Availability

All data needed to evaluate the conclusions in the paper are present in the paper and/or the Supplementary Materials. The RNA-seq data reported in this article has been deposited in NCBI’s Gene Expression Omnibus (GEO).

