## Supplemental figure for "Leptin Acts as a Peripheral Tropic Signal to Tune Steroidogenesis"

**This file includes:**

Figure. S1 to S8.

Table. S1 to S5.

Movie. S1 to S4.

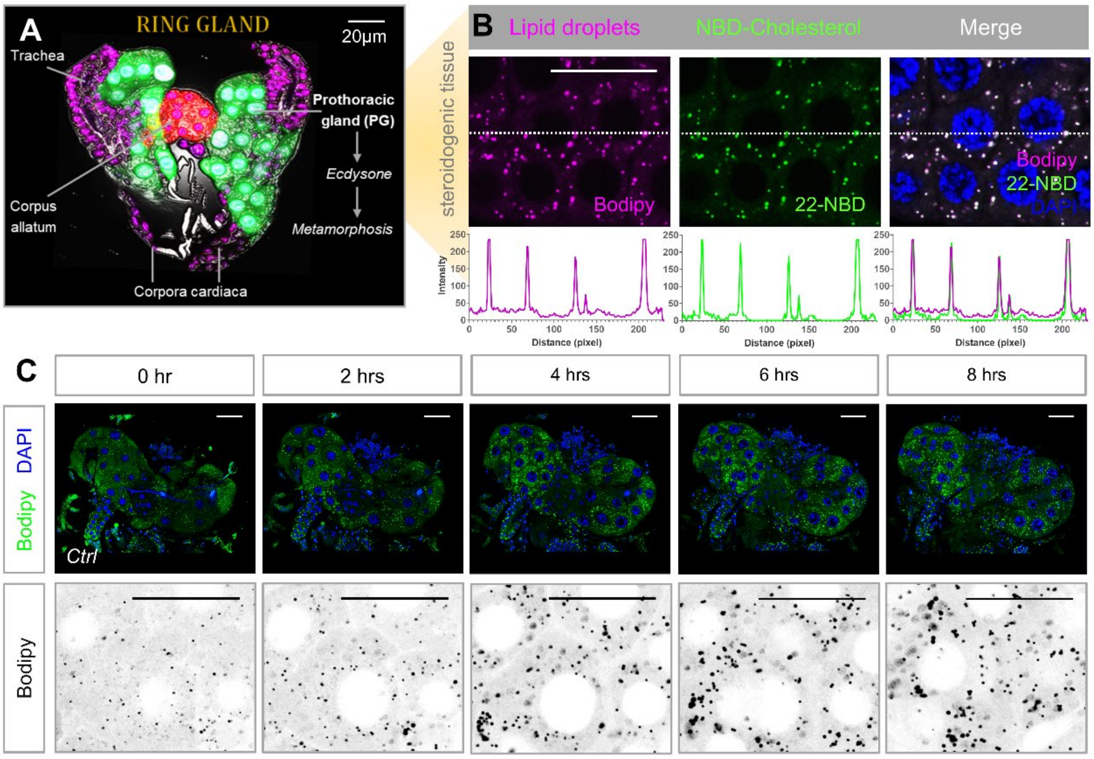


**Fig. S1 related Fig.1. LD and cholesterol dynamics in *Drosophila* steroidogenic tissues.** (A) A schematic of the ring gland, including the prothoracic gland, highlights its role in ecdysone production during metamorphosis. (B) Immunofluorescence staining reveals the presence of LDs (Bodipy, green) and cholesterol (22-NBD-Cholesterol, magenta) in steroidogenic tissues, with merged images showing co-localization. (C) Representative movie screenshots of real-time monitoring over 8 hours demonstrate the temporal dynamics of LD formation (Bodipy, green) with DAPI (blue) labling nuclei in control (*w1118*). Corresponding Bodipy-only images are displayed beneath each panel. Scale bars represent 20 μm for PG cells.

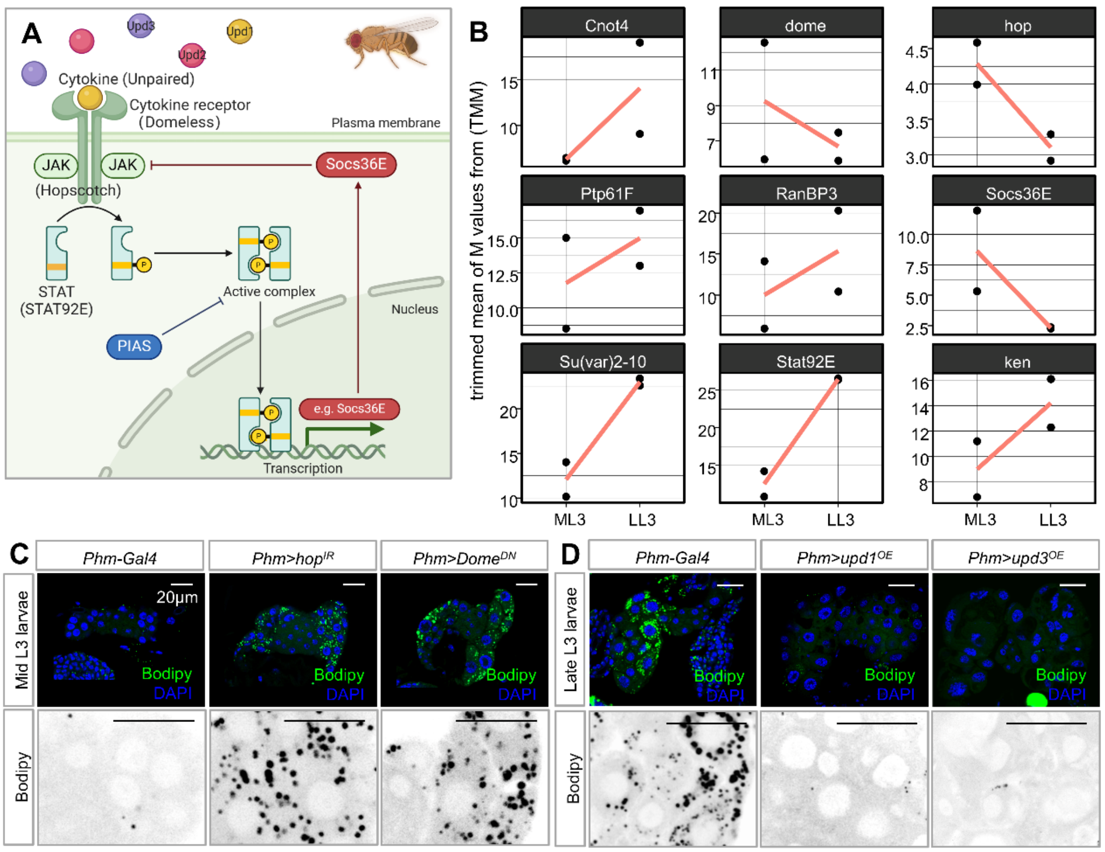


**Fig. S2 related Fig.2. Developmental expression profiles and functional validation of the JAK/STAT signaling pathway in the PG.** (A) Schematic diagram illustrating the components of the JAK/STAT signaling pathway in *Drosophila*. Unpaired cytokines (*upd1*, *upd2*, *upd3*) bind to the cytokine receptor *Domeless* (*dome*) on the plasma membrane, leading to activation of the Janus kinase *Hopscotch* (*hop*). Activated hop phosphorylates the transcription factor *Stat92E*, which forms an active complex and translocates to the nucleus to drive target gene transcription. Negative regulators include *Socs36E* (acting as a feedback inhibitor), *PIAS* (protein inhibitor of activated STAT, also known as *Su(var)2-10* in *Drosophila*), and other modulators such as *Ptp61F*, *RanBP3*, *Cnot4*, and *ken*. (B) Trimmed mean of M-values (TMM) normalized expression levels of JAK/STAT pathway genes in control (*w1118*) PG cells collected during the mid- and late- L3. Individual dots represent replicate samples, with lines connecting the group means to highlight diurnal changes in expression. Data derived from RNA-seq analysis. (C) Representative Bodipy (green) staining of LDs in the PG cell under control (*Phm-Gal4*), *hop RNAi* and *Dome* $^{DN}$ at mid-L3 stage. (D) At late-L3 larvae, representative images of LD size and density in the PG under control (*Phm-Gal4*) and animals with PG-specific overexpression of Upd1 or Upd3. Top panels show low-magnification confocal sections, and bottom panels present high-magnification views of representative regions indicated in the top panels.

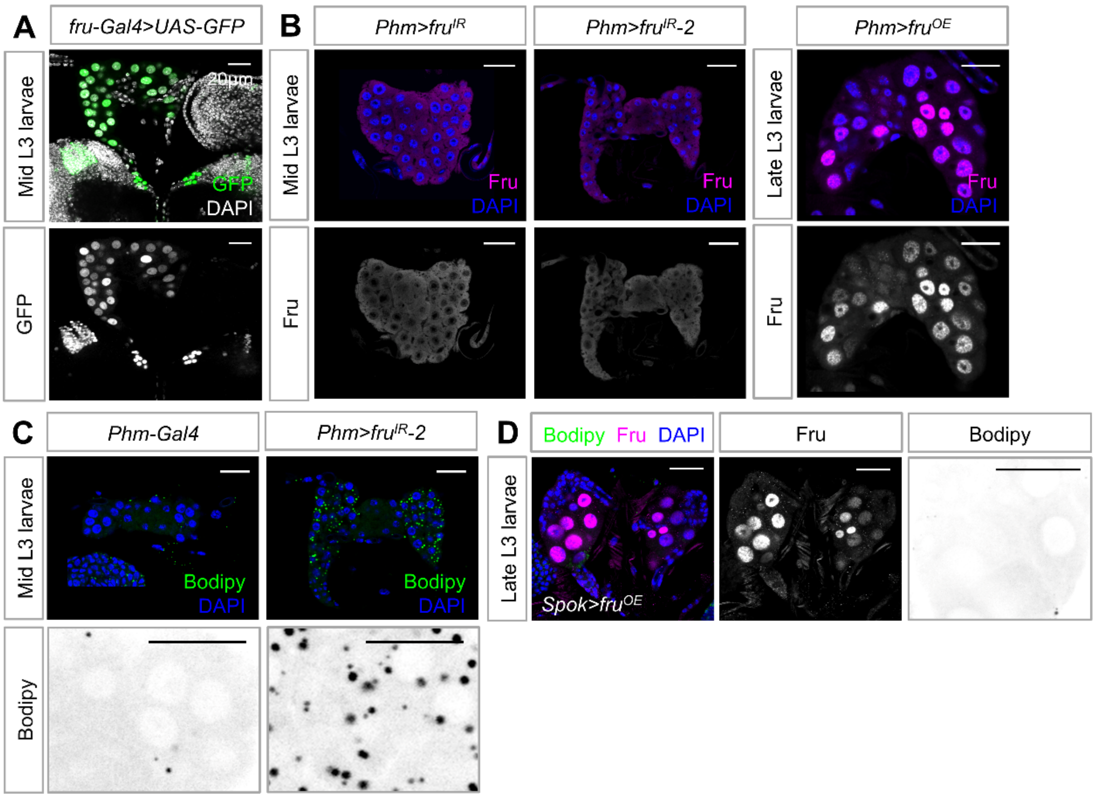


**Fig. S3 related Fig.3. Spatiotemporal Fru expression and functional validation in PG cells.** (A) Representative images showing the expression pattern of *UAS-GFP* driven by *fru-Gal4* in PG cells of mid-L3 instar larvae. (B) Anti-Fru antibodies (magenta) were used to stain PG cells with *fru* knockdown (all isoforms) at mid-L3 larvae (left) and with $Fru^{COM}$ overexpression at late L3 larvae (right). (C) Representative images of Bodipy (green) and DAPI (blue) in PG cells under control of *phm>* (PG-specific) with *fru* knockdown (*fru RNAi-2*) at mid-L3 larvae. Top panels show low-magnification confocal sections; bottom panels present high-magnification views of representative regions indicated in the top panels. (D) Representative images of Fru (magenta), Bodipy (green) and DAPI (blue) in PG cells under control of *Spok>* (PG-specific) with Fru overexpression at late L3 larvae. Scale bars, 20 μm in all confocal images.

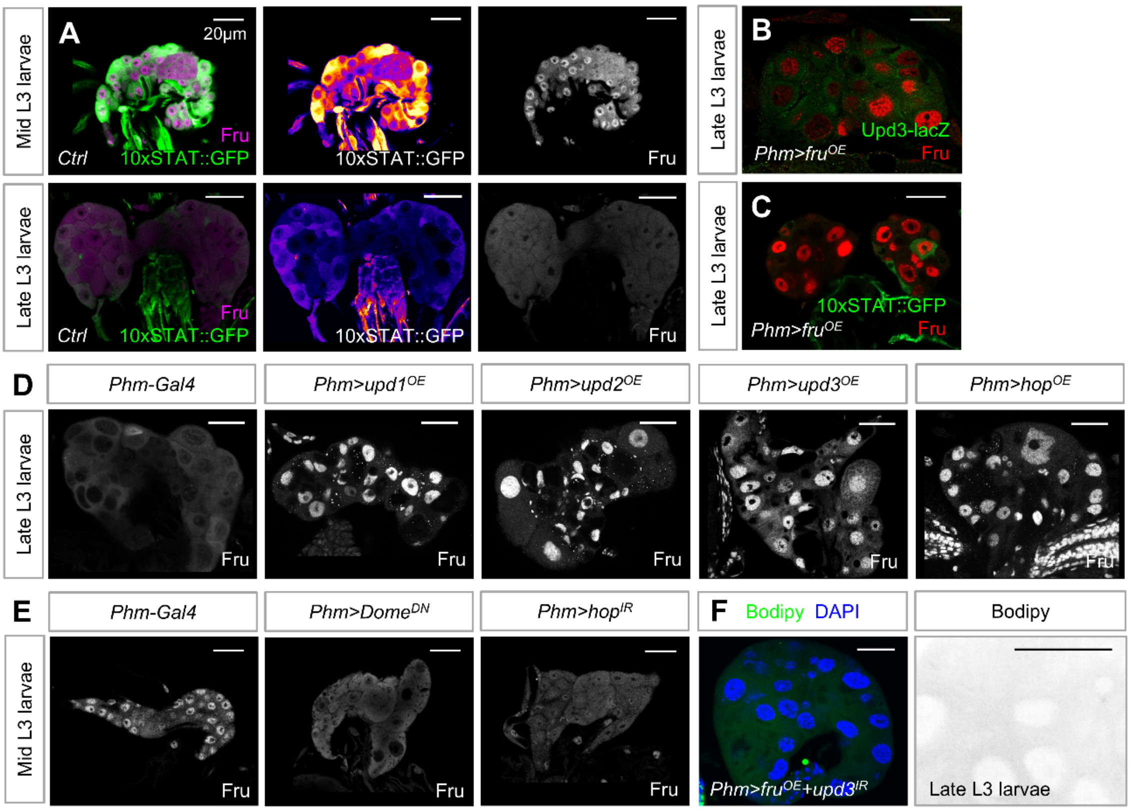


**Fig. S4 related Fig.3. Regulatory relationship between JAK/STAT signaling and Fru in the PG.** (A) Confocal images of the PG in JAK/STAT pathway activity reporter line (*10xSTAT::GFP*) showing co-localization of the Fru (magenta) with the GFP (green). Top panels: mid-L3 larvae; bottom panels: late L3 larvae. (B) Representative images showing the activity of the JAK/STAT pathway (*10xSTAT::GFP*, green, B) and the transcriptional level of *upd3* (*upd3-lacZ*, green, C) in the PG after PG-specific expression of Fru (red) in late L3 larvae. Representative images of Fru expression in PG cells under JAK/STAT activation (Upd1-3 and Hop overexpression) in late L3 larvae (D) and JAK/STAT inhibition (*Dome* and *Hop* knockdown) in mid-L3 larvae (E). (F) Representative images showing LD (Bodipy, green) in PG cells of late-L3 larvae with PG-specific *upd3* knockdown in Fru overexpression background. Scale bars, 20 μm in all channels.

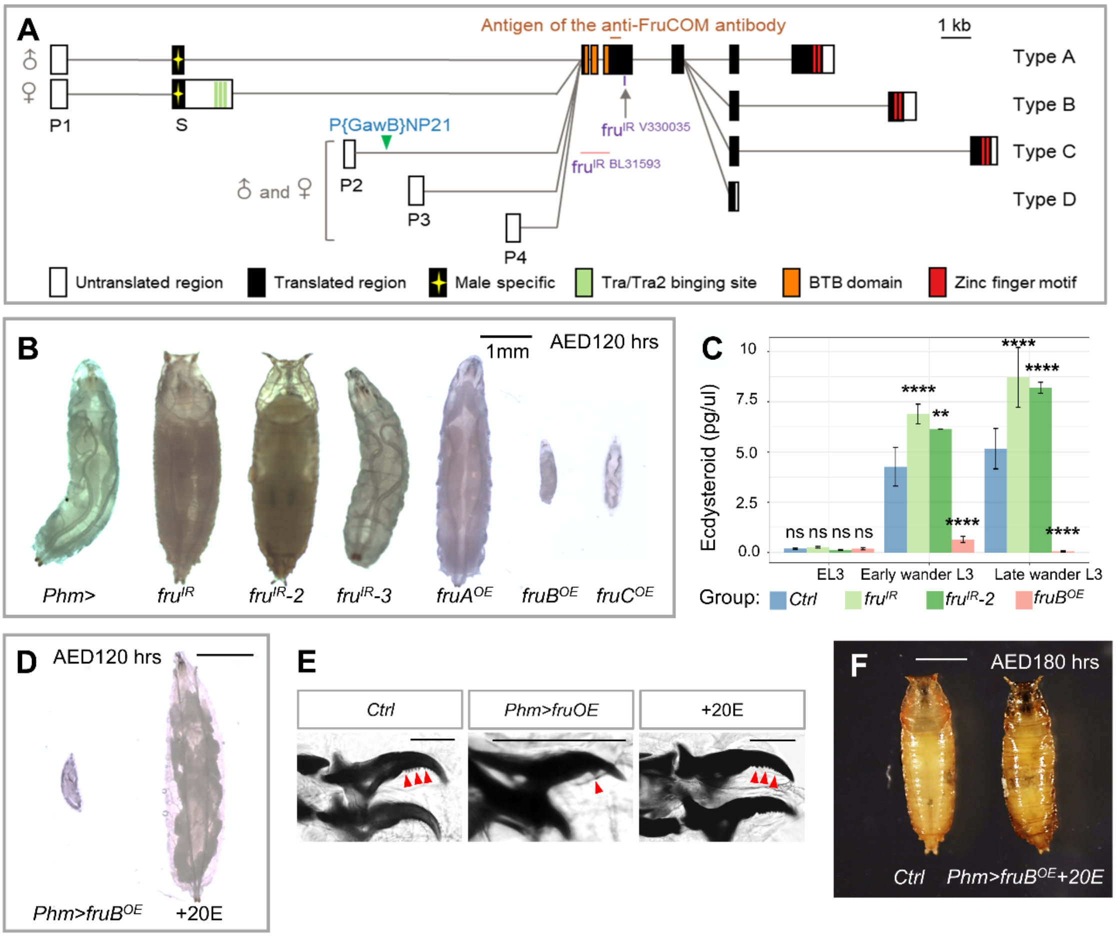


**Fig. S5 related Fig.3. Fru is essential for sexual developmental decision-making.** (A) Schematic representation of the *fru* gene locus. Locations of four promoters (P1-P4), the exon-intron organization, and the P-element insertion site of *fru*$^{NP21}$ (green triangles) are shown. Filled and open boxes indicate coding and non-coding exons, respectively. A-D denote isoform-specific exons for types A-D. The start and termination codons are also shown. The regions containing epitopes for the anti-FruCOM antibodies are indicated. (B) Larvae and prepupal phenotypes at 120 h after egg laying, including size defects and malformations. From left to right: control (*phm>*), PG-specific knockdown of *fru* (*fru RNAi 1-3*) and PG-specific overexpression of Fru (*fru A, B* and *C* isoform). (C) ELISA analysis measuring whole-body ecdysone levels in control, *fru* knockdown, and Fru overexpression animals at early L3, early wandering L3, and late wandering L3 stages. (D) Rescue experiments of the *Phm>fruB*$^{OE}$ developmental delay phenotype at 120 AED by dietary supplementation with 20-hydroxyecdysone (+20E). (J) Mouth hook structures of *Drosophila* larvae at 120 h AED. Control (*phm-GAL4*) exhibit L3 mouth hooks, whereas PG-specific overexpression of Fru results in L1 mouth hooks. Dietary supplementation with 20E rescues *Phm>fruB*$^{OE}$ larvae to L3 mouth hooks. (F) 20E treatment allows *Phm>fruB*$^{OE}$ animals to successfully pupariate by

180 h AED. Data are presented as mean ± s.e.m. ****p < 0.0001, ns, not significant. Scale bars represent 20 μm (PG), 1 mm (larvae) and 100 μm (mouth hooks).

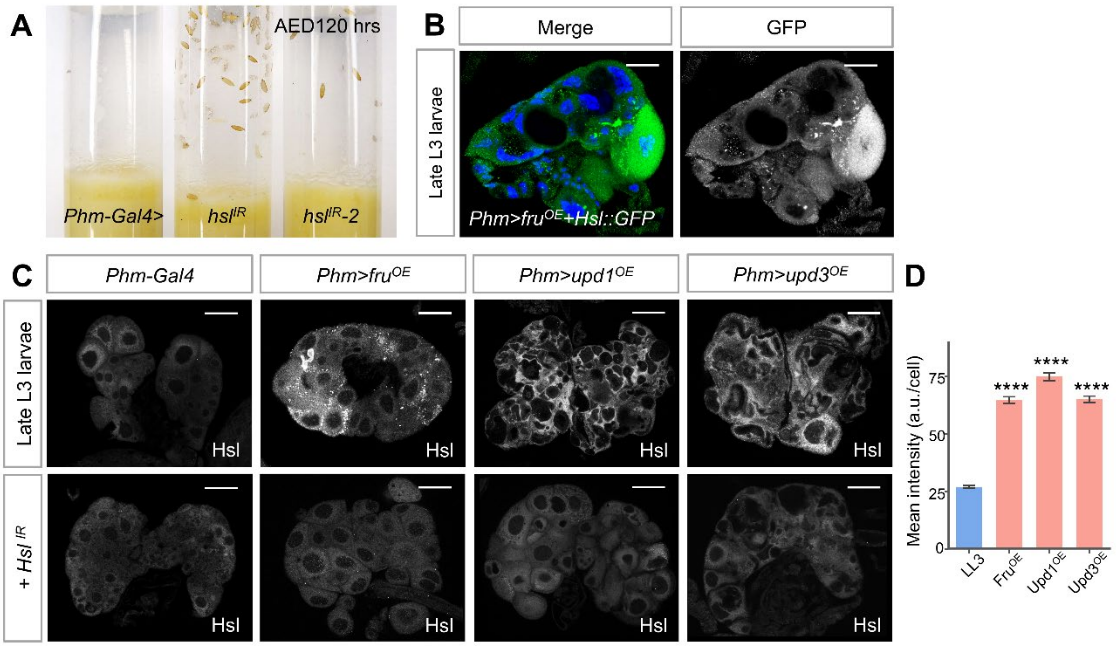


**Fig. S6 related Fig.4. JAK/STAT-Fru signaling regulates Hsl levels in the PG.** (A) Pupariation assay 120 hours AED showing pupariation status of larvae with PG-specific driven *hsl* knockdown using 2 independent RNAi lines (*Phm>hsl*$^{IR}$, *Phm>hsl*$^{IR}$*-2*) compared to controls (*Phm-Gal4*). (B) Representative confocal images of Hsl::GFP with DAPI (blue) in PG-specific Fru overexpression at late L3 larvae. (C) In late L3 larvae, immunofluorescence images showing Hsl expression in the PG under control, Fru overexpression, and JAK/STAT activation (from left to right: *Phm-Gal4, Phm>fru*$^{OE}$, *upd1*$^{OE}$ and *upd3*$^{OE}$) in the top panels, with corresponding *Hsl* knockdown shown in the bottom panels. (D) Quantitative analysis of Hsl fluorescence intensity in the PG for the genotypes shown in (C). Data are presented as mean ± s.e.m. ****p < 0.0001 (one-way ANOVA with multiple comparisons). Scale bars, 20 μm.

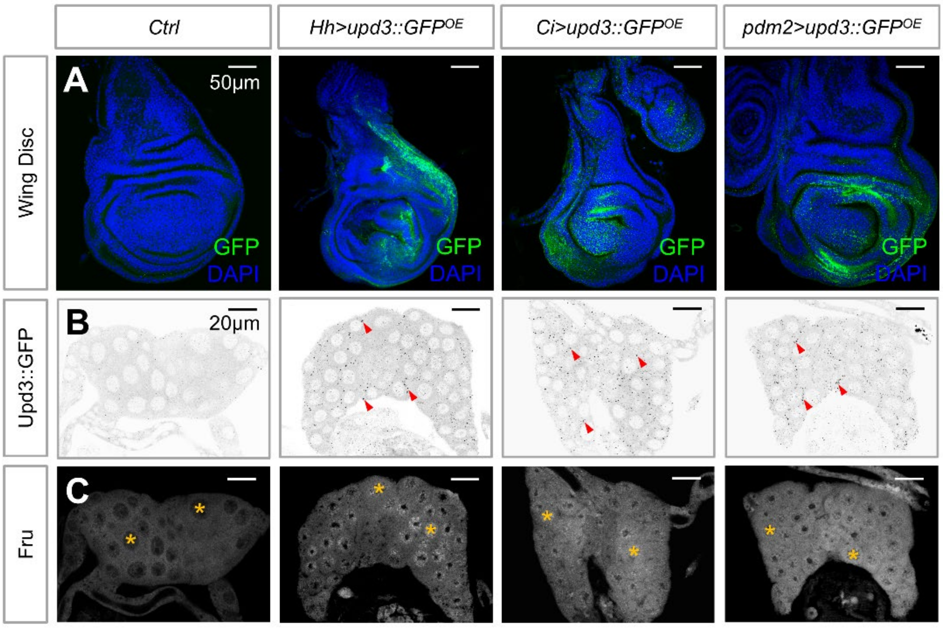


**Fig. S7 related Fig.6. Imaginal disc-derived Upd3 remotely modulates Fru expression in the PG**. Representative images showing the distribution of GFP signal in the wing imaginal disc (A) and the PG (B) following imaginal disc-specific overexpression of *upd3-GFP*. (C) Grayscale images representation of Fru expression levels in the PG corresponding to panel B. Red arrows indicate GFP puncta, and yellow asterisks denote nuclear Fru localization. Scale bars represent 20 µm for PG cells and 50 µm for wing imaginal disc.

**Table S1. Fly stocks and resources**

| Fly stocks | Source |
|---|---|
| *w1118* | BDSC#5905 |
| *phm-Gal4* | BDSC#80577 |
| *10×STAT::GFP* | [1] |
| *UAS-Stat92E RNAi* | VDRC #106980 |
| *UAS-3HA-STAT92E*$^{\Delta N\Delta C}$ | [2] |
| *UAS-hop RNAi* | BDSC#31319 |
| *UAS-Dome*$^{DN}$ | [3] |
| *UAS-upd1* | [4] |
| *UAS-upd3* | [5] |
| *UAS-fru RNAi* | BDSC#31593 |
| *UAS-fru RNAi-2* | VDRC#330035 |
| *UAS-fru RNAi-3* | VDRC#105005 |
| *UAS-upd2* | [6] |
| *UAS-hop* | BDSC#79033 |
| *Upd3-lacZ* | [7] |
| *Spok-Gal4* | BDSC#80578 |
| *UAS-GFP* | BDSC#5413 |
| *fru-NP21-Gal4* | BDSC#30027 |
| *UAS-Hsl RNAi* | BDSC#65148 |
| *UAS-Hsl RNAi-2* | VDRC#109336 |
| *UAS-Hsl* | [8] |
| *Hsl::GFP* | VDRC#318217 |
| *UAS-Soat1* | This study. |
| *UAS-upd2 RNAi* | BDSC#33988 |
| *UAS-upd3 RNAi* | BDSC#32859 |
| *30A-Gal4* | BDSC#37534 |
| *upd2*$^{\Delta}$*, upd3*$^{\Delta}$ | BDSC#55729 |
| *Act-Gal4/CyO* | BDSC#4414 |
| *UAS-NICD* | [9] |
| *hh-Gal4* | BDSC#600186 |
| *UAS-scribble RNAi* | VDRC #105412 |
| *UAS-lgl RNAi* | VDRC #51247 |
| *upd2_CBM-GFP* | [10] |
| *UAS-upd3::GFP* | [5] |
| *ci-Gal4* | [11] |

| | |
|---|---|
| *pdm2-Gal4* | BDSC#49828 |
| *UAS-fruC* | [12] |
| *Snr1 RNAi* | BDSC#32372 |
| *UAS-P35* | BDSC#5072 |

**Table S2. Primer sequences for RT-qPCR**

| **Primers** | Reference |
|---|---|
| GAPDH-F: TAAATTCGACTCGACTCACGGT | [13] |
| GAPDH-R: CTCCACCACATACTCGGCTC | [13] |
| upd2-F: CGGAACATCACGATGAGCGAAT | [14] |
| upd2-R: TCGGCAGGAACTTGTACTCG | [14] |
| upd3-F: ATCCCACCAATCCCCTGAAG | [15] |
| upd3-R: AGATTGCAGGTGTTCTCCCA | [15] |
| hLeptin-F: ACCCTGTGCGGATTCTTGTGG | [16] |
| hLeptin-R: CTCTGTGGAGTAGCCTGAAGC | [16] |
| hGAPDH-F: GGAGTCAACGGATTTGGT | [16] |
| hGAPDH-R: GTGATGGGATTTCCATTGAT | [16] |

**Other supplementary material for this manuscript includes the following:**

Table. S1 Fly stocks and resources.

Table. S2 Primer sequences for RT-qPCR.

Table. S3 Fluorescence intensity index analysis.

Table. S4 Quantification of lipid droplet area.

Table. S5 Statistics of RT-qPCR analysis.

Movie. S1 8-hour live monitoring the dynamics of lipid droplet formation (Bodipy, green) with nuclear staining (DAPI, blue) in the control PG (*w1118*).

Movie. S2 Enlarged grayscale images showing lipid droplet dynamics in the PG over an 8-hour live-imaging period, based on Movie S1.

Movie. S3 Live imaging of PG cells in 10xSTAT::GFP larvae over a 6-hour period, simultaneously capturing lipid droplet dynamics.

Movie. S4 LUT images show changes in JAK/STAT pathway activity during 6 hours of live imaging, based on Movie S3.
